# A Wnt-induced conformational phospho-switch in DVL3 controls interaction with Frizzled receptors and Wnt/β-catenin signaling

**DOI:** 10.1101/2025.09.10.675378

**Authors:** Miroslav Micka, Jitender Kumar, Petra Paclíková, Zuzana Hayek, Kateřina Hanáková, Cherine Bechara, Hana Plešingerová, Ondrej Šedo, Sara Bologna, Elise Del Nero, Kristína Gömöryová, Vojtěch Bystrý, Tomáš Gybeľ, Tereza Číhalová, Marek Kravec, David Potěšil, Zbyněk Zdráhal, Konstantinos Tripsianes, Vítězslav Bryja

## Abstract

When Wnt ligands bind to Frizzled (FZD) receptors, Dishevelled protein (DVL) gets multiphosphorylated by Casein kinase 1 (CK1). Although, it is well known that DVL phosphorylation relays Wnt signals from receptors to downstream effectors, any mechanistic aspects of DVL function related to phosphorylation remain unresolved. Here, we uncovered a Wnt-induced DVL phospho-switch which is mutually exclusive with FZD engagement. CK1 multiphosphorylation changes dramatically the bulk electrostatics to promote DVL intramolecular coupling between the DEP domain and the adjacent disordered region. A panel of phospho-switch mutants demonstrated a switch-like coupling at the molecular level when a charge threshold is reached. Charge accumulation proximal to DEP proved to be a key functional event required, but not sufficient, for Wnt/β-catenin signaling. Interestingly, the charge-dependent switch-like character of DVL controlled its association with FZD. By integrating findings at different levels, we propose a universal mechanism in which Wnt-induced DVL conformational phospho-switch outcompetes FZD binding and triggers DVL detachment from FZD.

## Introduction

Wnt signaling is an essential developmental pathway that is highly conserved in all metazoans. It orchestrates embryonic development and maintains adult tissue homeostasis. At the cellular level, activation of Wnt signaling triggers responses such as mitotic activity, differentiation, and the establishment of polarity. However, aberrant activation of the Wnt pathway can lead to various types of cancer and other diseases.^1^

The central component of Wnt signaling is the cytoplasmic protein Dishevelled (DVL), which has three paralogs (DVL1, DVL2, and DVL3) in mammals. DVL has a scaffolding role and is critical for signal transduction in Wnt pathways. Genetic evidence of DVL’s requirement exists in numerous model organisms. For example, in mice, DVL knockout (KO) leads to developmental defects, which are aggravated by the KO of multiple DVL hosmologs.^2–4^ . The critical importance of DVL for the Wnt/β-catenin signaling is demonstrated by the CRISPR KO of all three DVL homologs also in cells, which are not able to transduce the Wnt signal.^5,6^

DVL is a modular protein made up of three globular domains (DIX, PDZ, DEP) connected by intrinsically disordered regions and a long C-terminal disordered tail (IDRs). The current knowledge about its molecular functions has been built up through piecemeal experimental investigation of the well-structured domains^5,7–12^ while the disordered regions, that account for 2/3 of the DVL sequence, have yet to be explored in structural and functional investigations. Generally, the IDRs, including those of DVL, are rich in phosphorylation sites,^13^ suggesting that DVL mechanistic secrets reside also in the disordered regions and most likely in their synergistic function with the structured domains.

The current model of DVL function in the Wnt/β-catenin pathway suggests that upon binding of Wnt to its receptor Frizzled (FZD), DVL is recruited to the plasma membrane, where it directly interacts with FZD through its DEP domain^7,14^ and oligomerizes. An essential feature of DVL in Wnt/β-catenin signaling is the formation of homopolymers or heteropolymers with Axin mediated through their DIX domains.^8^ Recruitment of Axin to the membrane and specifically to phosphorylated LRP6 leads then to the inhibition of the β-catenin destruction complex followed by β-catenin stabilization and downstream signaling.^9,15–19^

In response to Wnt ligands, endogenous DVL becomes phosphorylated by Casein kinase 1 (CK1) δ or ε. The massive phosphorylation of DVL is manifested as a mobility shift on SDS-PAGE, making it a simple readout.^20–22^ Several studies have mapped the DVL phosphorylations mediated by CK1 ^23–29^. Mutation of some of the phosphosites to alanine had no effect on DVL’s role in Wnt/β-catenin signal transduction,^24,27,28^ while mutation of other sites impaired DVL’s ability to activate Wnt/β-catenin.^23,25^ However, none of these phosphorylation sites proved to be truly essential for β-catenin activation and their mutation never resulted in a completely nonfunctional DVL. Most of these studies used experimental systems with overexpressed DVL for both detection of phosphorylation as well as subsequent functional testing,^25,26,28,29^ which could limit the conclusions.

Here, we sought to identify endogenous DVL3 phosphorylation sites induced by Wnt ligands under physiological conditions and elucidate their consequences on DVL function and signaling. Our analysis pointed to an S/T cluster of IDR2 proximal to the DEP domain that is heavily phosphorylated by CK1ε. The cumulative electrostatic interactions drive a switch-like response and couple the phosphorylated IDR2 with DEP. This conformational change is incompatible with DEP binding to FZD receptors. Comparative proteomics revealed FZDs as the prominent effectors of the DVL phospho-switch function providing a direct link to the mechanistic findings. Our data support that IDR2 phosphorylation, followed by DVL3 conformational coupling is an essential step required for DVL to dissociate from FZDs and to signal towards downstream effectors.

## Results

### Wnts induce phosphorylation of endogenous hDVL3 at S394

In order to investigate the Wnt-induced changes in DVL phosphorylation at the endogenous level, we treated FreeStyle™ 293-F cells with Wnt-3a, Wnt-5a, or control conditioned medium (CM). Cells were pre-treated with the Porcupine inhibitor LGK974 to block autocrine Wnt production and to reset the system to a completely OFF state. Subsequently, endogenous DVL3 was purified using a two-step protocol. We took advantage of a naturally occurring cluster of histidine (His) residues at the C-terminus of DVL3 and performed a His pulldown, followed by immunoprecipitation (IP) with an anti-DVL3 antibody. Next, the samples were resolved on SDS-PAGE (**Fig. S1**) and the bands corresponding to DVL3 were analyzed by MS (**Fig. 1A**). Numerous phosphorylated residues were detected; however, only phosphorylation of a highly conserved serine (S394) in the second disordered region (IDR2) of hDVL3 was significantly enriched after Wnt treatment (**Fig. 1B**). Interestingly, S394 phosphorylation of endogenous DVL3 was the result of both Wnt-3a and Wnt-5a stimulation (**Fig. 1B, 1C, Fig. S2**).

**Figure 1.**
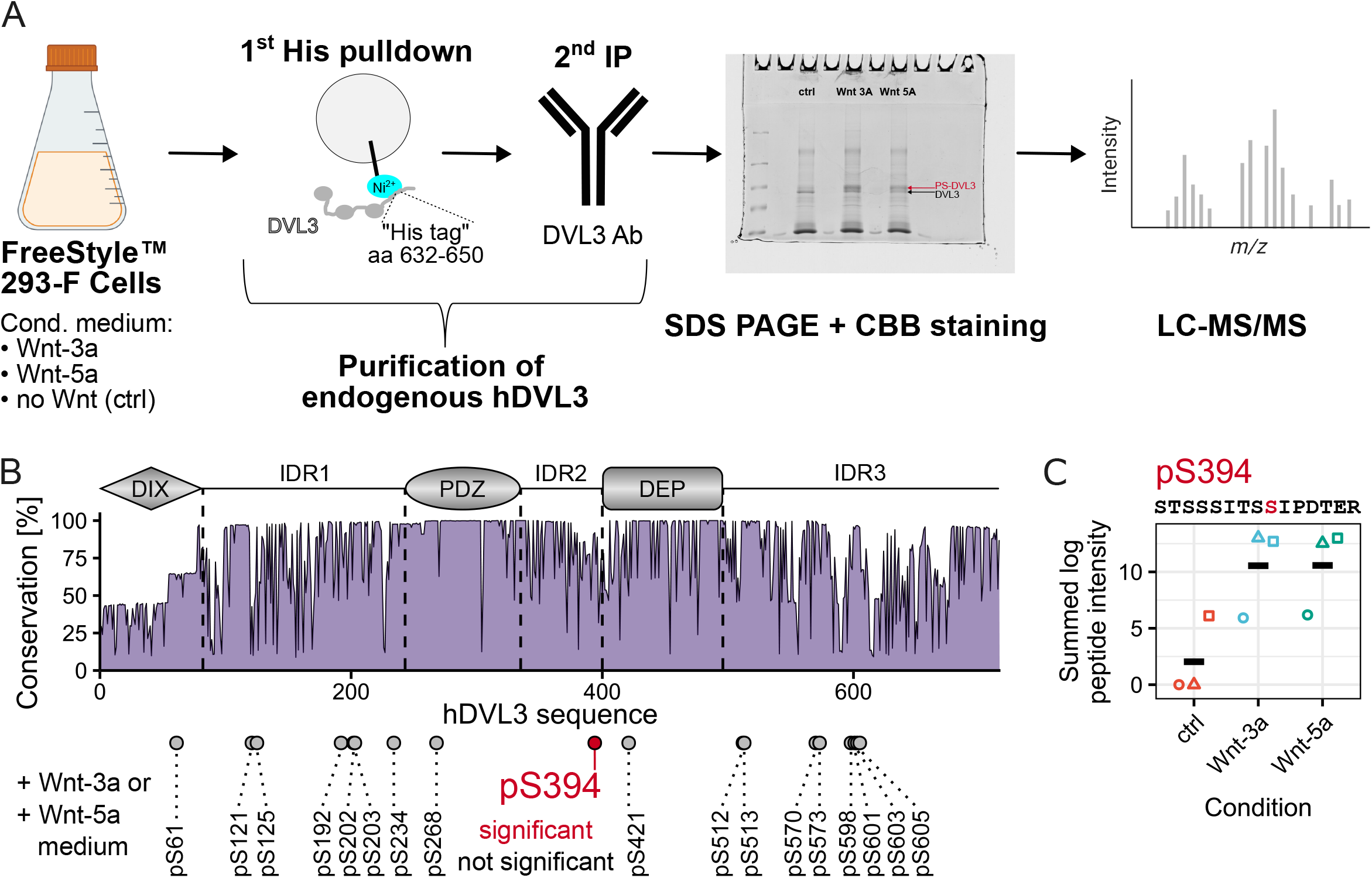
Wnts induce phosphorylation of endogenous hDVL3 at S394. **A**. Schematic workflow to detect Wnt-induced phosphorylations of endogenous hDVL3. **B**. Amino acid conservation across the hDVL3 sequence from a multiple sequence alignment of 854 DVL sequences. The hDVL3 architecture is shown at the top and the detected phosphorylation sites are depicted as circles at their respective positions at the bottom. Phosphorylation site S394 is highlighted in red due to its statistical significance compared to the control conditioned medium (without Wnts) experiment. Statistical significance was tested by ANOVA at 90% level of confidence. N = 3. **C**. Plot of the S394 phosphorylation site in endogenous hDVL3. Different shapes of points represent different replicates. The mean values are indicated by black dashes. The sequence of the peptide where the phosphorylation of S394 was detected by LC-MS/MS is shown at the top, S394 is highlighted in red.

### CK1δ and CK1ε efficiently catalyze multisite phosphorylation of DVL3 IDR2 *in vitro*

S394 is part of the disordered region (IDR2: aa 335-396) between the PDZ and DEP domains of DVL3. IDR2 is relatively conserved among 854 DVL homologs across hundreds of animal species (**Fig. 1B**) and in human paralogs displays a common arrangement of multiple S/T sites organized in three clusters (**Fig. 2A**). We have hypothesized that IDR2 S/Ts are phosphorylated by the well-described DVL kinase – CK1ε – that mediates the typical Wnt-induced phosphorylation-dependent DVL3 mobility shift (**Fig. 2B**).^21,22,30–32^ To confirm whether IDR2 and S394 in particular is a primary target of CK1ε, we purified hDVL3 IDR2 and phosphorylated it *in vitro*. The gradual mobility shift of IDR2, corresponding to increasing levels of phosphorylation suggested multisite phosphorylation by CK1ε (**Fig. 2C)**. Intact mass analysis by matrix assisted laser desorption ionization (MALDI)-MS confirmed a time-dependent and quantitative accumulation of multiple phosphorylated IDR2 species; on average, 11 phosphate groups were added to a single IDR2 molecule, with a maximum of 15 detected at the end of the reaction (**Fig. 2D, Fig. S3A**). Paradoxically, earlier LC-MS/MS analysis of full-length DVL3 phosphorylation showed that IDR2 is phosphorylated poorly or not at all.^29^ This is likely due to technical limitations of LC-MS/MS in detecting peptides phosphorylated at more than 3-4 sites.^33,34^ We demonstrated experimentally that these limitations apply as well to the *in vitro* multisite phosphorylated IDR2 peptides, which could be detected once partially dephosphorylated (**Fig. S3B)**. Of note, this validation also shows that LC-MS/MS can still be useful in the indirect quantification of multisite phosphorylation by monitoring the decrease in the intensity of the non-modified species during the kinase reaction.

**Figure 2.**
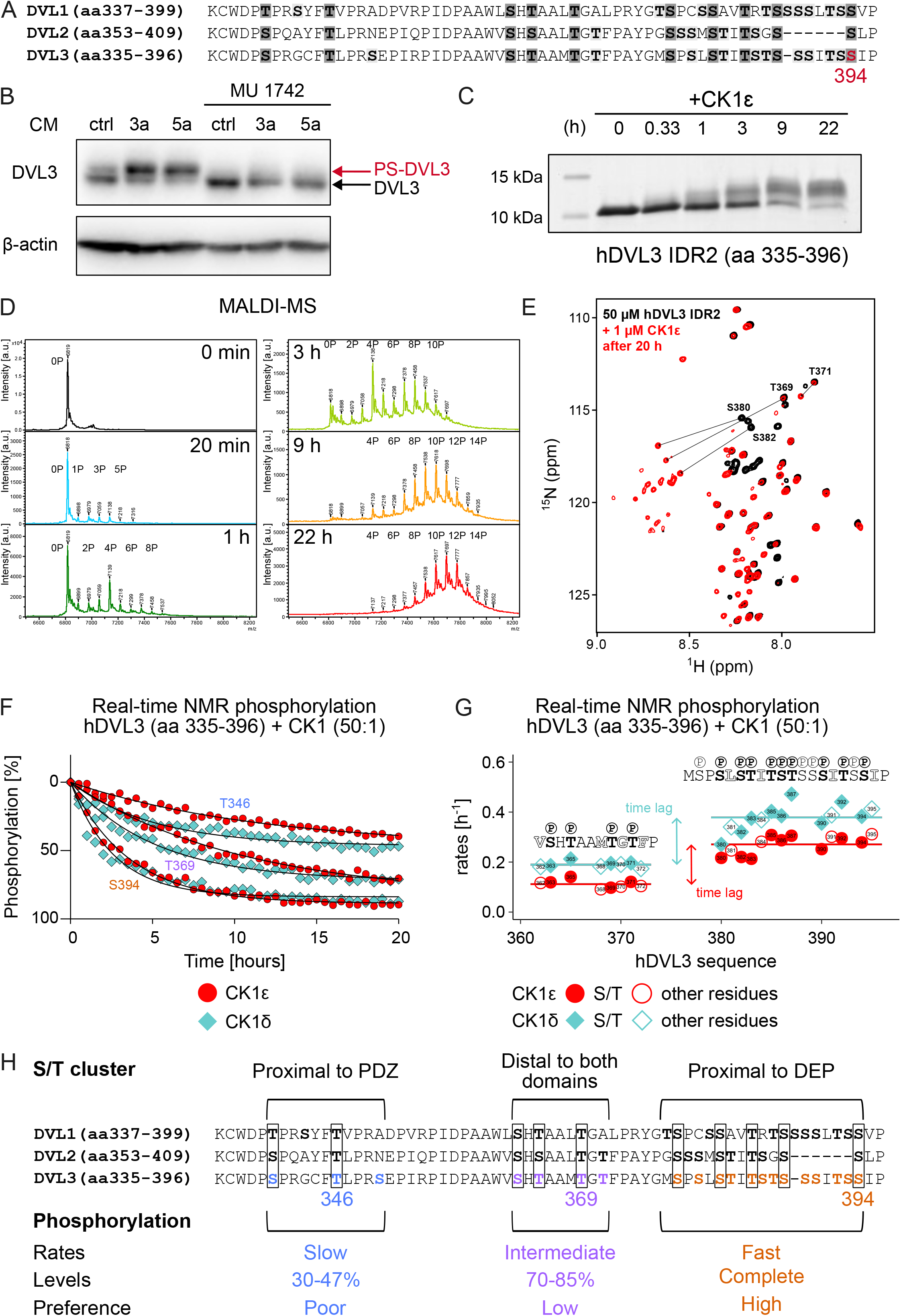
*In vitro* multisite phosphorylation of hDVL3 IDR2 by CK1δ and CK1ε. **A**. Conservation of IDR2 among all three human DVL homologs. Conserved S/T residues are highlighted in dark grey, other S/T residues are highlighted in light grey. S394 of hDVL3 is highlighted in red. **B**. CK1ε inhibition by 10 µM MU1742 attenuates the DVL3 mobility shift on SDS-PAGE induced by Wnt conditioned media (CM). β-actin was used as the loading control. LGK974 (0.2 µM) was applied to block autocrine Wnt stimulation. **C**. Time course of *in vitro* phosphorylation of hDVL3 IDR2 (aa 335-396) by CK1ε. 50 µM hDVL3 IDR2 was phosphorylated by 1 µM CK1ε in the presence of 10 mM MgCl_2_ and 2 mM ATP. **D**. MALDI-time of flight (TOF) mass spectra of ^15^N-labelled hDVL3 IDR2 over the time course of *in* vitro phosphorylation by CK1ε. IDR2 phosphorylated species are annotated with numbers **E**. Overlay of ^1^H-^15^N HSQC spectra of hDVL3 IDR2 (black) and hDVL3 IDR2 phosphorylated *in vitro* by CK1ε (red). Phosphorylation-induced changes in the chemical shift of S/T residues are highlighted by black arrows. **F**. Phosphorylation kinetics analyzed by real-time NMR for 20 hours as the decrease in peak intensity of S/T residues. T346, T369 and S394 are representatives of the S/T clusters proximal to the PDZ domain, distal to both domains, and proximal to the DEP domain, respectively. Time points of phosphorylation by CK1ε are depicted as red circles, time points of phosphorylation by CK1δ are depicted as cyan diamonds. **G**. Phosphorylation rates for individual S/T or neighboring residues of hDVL3 IDR2 for the distal and proximal S/T clusters to DEP. Solid lines represent mean phosphorylation rates for the cluster with fast (0.27 hours^-1^, 0.38 hours^-1^) and intermediate (0.11 hours^-1^, 0.19 hours^-1^) kinetics for CK1ε (red) or CK1δ (cyan) phosphorylation, respectively. S/T residues are represented by full-color symbols, neighboring residues are represented by outlines. **H**. Summary of DVL3 phosphorylation kinetics with respect to IDR2 S/T clusters projected onto the sequences of the three human DVL paralogs.

To obtain quantitative, residue-specific information and map unambiguously the DVL3 phosphosites targeted by CK1ε, we performed time-resolved NMR experiments. Phosphorylation of S/T residues is easily recognizable, since their peak resonances are downfield shifted outside the original spectrum (**Fig. 2E)**. Using the continuous NMR readout, the phosphorylation kinetics of every S/T residue were accessible as the decrease in peak intensity of the non-phosphorylated IDR2 (**Fig. 2F and Fig. S4**). Non-phosphorylatable residues adjacent to S/T sites were also included in the analysis to increase the profiling accuracy of CK1ε phosphorylation (**Fig. S5**). The phosphorylation events, as determined by the pseudorates and occupancy levels per S/T site, differed markedly between the three S/T clusters of IDR2. The S/T cluster proximal to the DEP domain exhibited the fastest kinetics and complete phosphorylation (exemplified by S394), the central S/T cluster distal to both DEP and PDZ, showed slower kinetics and an extent of phosphorylation between 70-85% (exemplified by T369), and the S/T cluster proximal to the PDZ domain was poorly phosphorylated reaching levels between 30-47% (exemplified by T346) (**Fig. 2F, 2H Fig. S4**). Remarkably, IDR2 phosphorylation by the closely related and functionally redundant^22,35^ CK1δ produced nearly identical kinetic parameters (Pearson’s correlation coefficient of phosphorylation rates: 0.97) (**Fig. 2F, G** and **Fig. S3C, D**) and validated the cluster-based phosphorylation mechanism. Both phosphorylation datasets suggest that there is a high preference for the cluster proximal to DEP and only when it is saturated, then the distal cluster is phosphorylated generating a time lag in the phosphorylation of the two clusters (**Fig. 2G**).

### CK1ε-induced phosphorylation profile is faithfully reproduced across the order-disorder continuum of DVL3

To investigate CK1ε-mediated phosphorylation in a more physiological and functional context, we studied the IDR2 physically coupled to its flanking domains, PDZ and DEP (PDZ-IDR2-DEP: aa 243-496). We phosphorylated PDZ-IDR2-DEP *in vitro* by CK1ε for up to 12 hours (**Fig. 3A**). Similarly to the isolated IDR2, a gradual shift was observed on SDS-PAGE, with the phosphorylation levels increasing over time. Quantification of the phosphorylation at selected time points (0 h, 1 h, 4 h, 12 h) by LC-MS/MS confirmed a robust, time-dependent phosphorylation of IDR2 residues. The occupancy levels determined by LC-MS/MS (**Fig. 3A**, right) were in full agreement with the cluster-based phosphorylation mechanism deduced from the analysis of IDR2 alone (**Fig. 2**). Of note, phosphorylation of the high-preference cluster (aa 377-400), proximally to the DEP domain, was analyzed indirectly, based on the decrease of intensity of the non-phosphorylated peptide (see **Fig. S3B**). Importantly, no quantitative phosphorylation was detected for S/T residues residing in the PDZ or DEP domains. The intact mass analysis of PDZ-IDR2-DEP by native electrospray ionization (ESI)-MS (**Fig. 3B**) confirmed extensive phosphorylation and a mass increase corresponding to the addition of 8 to 19 phosphates, in line with the MALDI-MS analysis of IDR2 (**Fig. 2D**).

**Figure 3.**
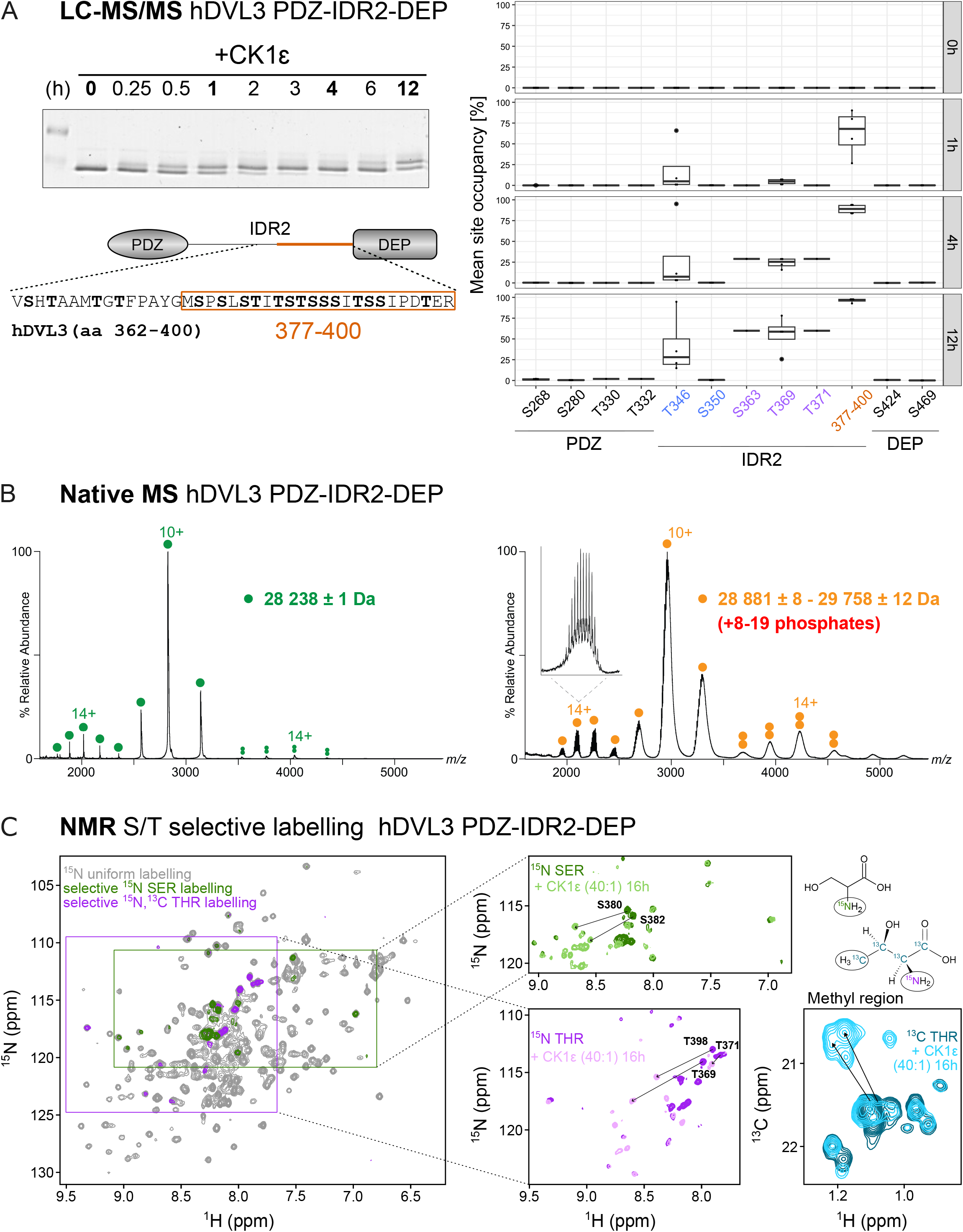
CK1ε-induced phosphorylation profile in PDZ-IDR2-DEP. **A**. Time course of *in vitro* phosphorylation of hDVL3 PDZ-IDR2-DEP (aa 243-496) by CK1ε. 5 µM hDVL3 PDZ-IDR2-DEP was phosphorylated by 125 nM CK1ε in the presence of 10 mM MgCl_2_ and 1 mM ATP. A representative gel used for the analysis with selected time points (in bold) analyzed by LC-MS/MS. The results are displayed as mean site occupancy. N = 4. Sites belonging to S/T clusters proximal to PDZ, distal to both domains, and proximal to DEP are highlighted in blue, purple, and orange, respectively **B**. Native ESI-MS spectra of PDZ-IDR2-DEP in the non-phosphorylated (left, green) and phosphorylated (right, orange) forms, showing the presence of two main conformations for the monomeric species: a more open and a more closed conformation with average charge state distribution of 14+ and 10+ respectively. Non-phosphorylated monomers were detected at 28,238 ± 1 Da, whereas multiple phosphorylated isoforms were detected between 28,881 ± 8 Da and 29,758 ± 12 Da, corresponding to 8-19 phosphates, resolved for the low m/z charge state distribution of the PDZ-pIDR2-DEP (the detailed segment). Homodimeric species were detected in both cases (double circles) at 56,521 ± 6.5 Da for the non-phosphorylated form, and an average 59,143 ± 115 Da for the phosphorylated one. **C**. Overlay of ^1^H-^15^N HSQC spectra of ^15^N-uniformly (grey), ^15^N-serine-selective (green), and ^15^N/^13^C-threonine-selective (magenta) labeled PDZ-IDR2-DEP. Insets show an overlay of spectra before and after phosphorylation *in vitro* by CK1ε. Arrows indicate phosphorylation-induced chemical shift changes of S/T residues.

Next, we employed NMR to study CK1ε multisite phosphorylation across the order-disorder continuum of DVL3. To analyze PDZ-IDR2-DEP phosphorylation in the very complex NMR spectra, we utilized ^15^N-serine-selective and ^15^N/^13^C-threonine-selective labeling. This approach reduces the number of NMR-visible signals to the ones concerned and enables unambiguous assignment of the spectral fingerprints of S/T signals (**Fig. 3C**). Indeed, all S/T residues (residing in PDZ, IDR2, or DEP) were identified in the non-phosphorylated form. CK1ε phosphorylation led to the characteristic chemical shift changes for IDR2 peaks, but no downfield changes were observed for the PDZ or DEP peaks. As an extra proof of threonine phosphorylation, we monitored the methyl probes by NMR. The covalent addition of a phosphate group to the hydroxyl moiety of threonine alters the chemical environment of the nuclei in its vicinity, leading to an upfield carbon chemical shift (**Fig. 3C**). These data confirm that the adjacent structured domains do not interfere with the CK1ε multisite phosphorylation mechanism of IDR2. Yet, this mechanism does not provide any functional cue in regard to DVL3 regulation or function.

### Multisite phosphorylation of IDR2 acts as a charge-driven conformational switch in DVL3

To gain mechanistic insights into Wnt-induced DVL3 phosphorylation, we obtained phosphorylated DVL3 proteins in high homogeneity and yield directly from *E. coli* cells by co-expression with CK1ε. A comparison between PDZ-IDR2-DEP and PDZ-pIDR2-DEP spectra revealed several chemical shift changes along with peak broadening (**Fig. 4Ai**). The effects of IDR2 phosphorylation were prominent on the DEP domain indicating a change in the conformation of PDZ-IDR2-DEP. In principle this can be either due to pIDR2 binding to DEP or by DEP swapping from monomeric to dimeric species, as reported previously.^7^

**Figure 4.**
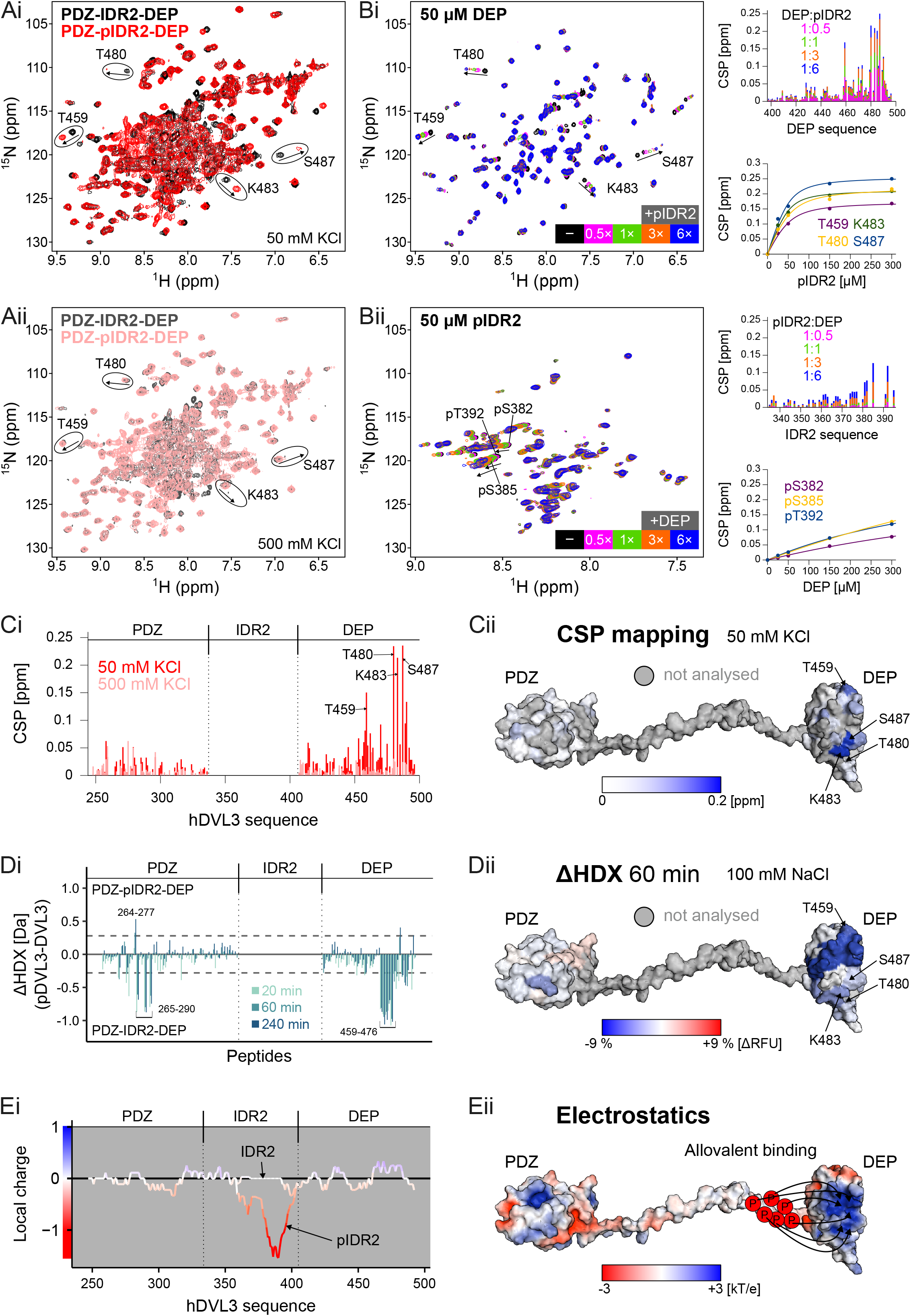
Multisite phosphorylation of IDR2 acts as a charge-driven conformational switch in DVL3. **Ai**. Overlay of ^1^H-^15^N HSQC spectra of hDVL3 PDZ-IDR2-DEP (aa 243-496, black) and phosphorylated PDZ-IDR2-DEP (PDZ-pIDR2-DEP, red) in the presence of 50 mM KCl. Arrows indicate chemical shift perturbations (CSPs) for selected DEP residues (circled) due to IDR2 phosphorylation. **Aii**. Overlay of ^1^H-^15^N HSQC spectra of hDVL3 PDZ-IDR2-DEP (dark grey) and PDZ-pIDR2-DEP (pink) in the presence of 500 mM KCl. Circles and arrows are used as reference points to contrast the lack of phosphorylation-induced CSPs in high salt. **Bi**. Overlay of ^1^H-^15^N HSQC spectra of ^15^N-labeled 50 µM hDVL3 DEP domain free and in the presence of increasing amounts of phosphorylated hDVL3 IDR2 (pIDR2). Arrows indicate CSPs for the selected residues. CSPs throughout the DEP sequence and binding isotherms for the indicated residues are plotted. **Bii**. Overlay of ^1^H-^15^N HSQC spectra of ^15^N-labeled 50 µM hDVL3 pIDR2 free and in the presence of increasing amounts of hDVL3 DEP. Arrows indicate CSPs for selected phosphorylated residues. CSPs throughout the IDR2 sequence and binding isotherms for the indicated residues are plotted. **Ci**. Phosphorylation-induced CSPs throughout the PDZ-IDR2-DEP sequence in the presence of 50 mM KCl (red) or 500 mM KCl (pink). **Cii**. Phosphorylation-induced CSPs in the presence of 50 mM KCl mapped onto the PDZ-IDR2-DEP structure. CSP magnitude is color-coded, ranging from white (0 ppm) to blue (0.2 ppm). Residues that could not be analyzed are colored grey. **Di**. Differential relative deuterium uptake (ΔHDX, phosphorylated – non-phosphorylated PDZ-IDR2-DEP) in Da for detected peptides in the PDZ and DEP domains at 20 min, 60 min, and 240 min. The dotted line indicates the 99% confidence interval as a threshold for significance at 0.28 Da, calculated using the Hybrid statistical test. **Dii**. Percentage of differential relative fractional uptake (ΔRFU) of the HDX-MS experiment mapped onto the PDZ-IDR2-DEP structure at 60 minutes time point. Phosphorylation-induced reduction in deuterium uptake is represented in blue, while increase in deuterium uptake is in red, according to the scale. Residues not analyzed are colored grey. **Ei**. Local charge distribution of PDZ-IDR2-DEP and PDZ-pIDR2-DEP calculated at pH=6.5. **Eii**. Electrostatic surface of PDZ-IDR2-DEP with negative and positive charges colored red and blue, respectively. Arrows indicate interaction of negatively charged phosphate groups in IDR2 with the positively charged cleft of the DEP domain.

To distinguish between the two mechanistic possibilities, and to gain more insights into the role of IDR2 phosphorylation, we disengaged order from disorder and performed experiments using the individual components. PDZ domain, DEP domain, IDR2, and pIDR2 were differentially labeled and the potential interactions between order and disorder were studied using an NMR-based binary approach. We performed NMR titrations by addition of excess IDR2 to the PDZ or DEP domain, and *vice versa*, by addition of excess PDZ or excess DEP domain to IDR2. None of the combinations yielded chemical shift perturbations (CSPs) to suggest a physical interaction (**Fig. S6A, B**). Next, we performed the same titrations using pIDR2. IDR2 phosphorylation did not yield spectral changes when combined with PDZ (**Fig. S6C**). On the contrary, serial addition of pIDR2 induced large CSPs in the DEP domain (**Fig. 4Bi**). On saturation, the linear trajectory of the DEP shifting resonances coincided with the peak positions in PDZ-pIDR2-DEP. The CSPs quantified *in trans* and the CSPs induced by phosphorylation of IDR2 *in cis* were nearly identical and pinpointed the same binding interface when mapped onto the DEP structure (**Fig. 4Bi, 4C, S6D**).

The specific interaction between DEP and pIDR2 was corroborated by monitoring the chemical shift changes in pIDR2 spectra as well (**Fig. 4Bii**). Although we could only assign a fraction of the phosphorylated S/T peaks in pIDR2, pronounced CSP effects *in trans* were detected for the phosphorylated peaks, which in native context (PDZ-IDR2-DEP) are part of the DEP proximal cluster, suggesting that this is the primary binding site with DEP when IDR2 is phosphorylated by CK1ε. However, when the NMR binding isotherms of pIDR2 were compared to the NMR binding isotherms of DEP, we noticed a striking difference, though the two titrations were performed identically to analyze the same interaction. DEP isotherms were saturated, whereas pIDR2 isotherms were not. This suggests a dynamic interaction between a single binding site mapped on the ordered domain (DEP) and any of the multiple phosphate groups present in the disordered region (pIDR2), a phenomenon known as allovalency.^36,37^

In order to validate the intramolecular communication between pIDR2 and DEP, we performed differential Hydrogen/Deuterium exchange coupled to mass spectrometry (HDX-MS) experiments. We compared the deuteration uptake of PDZ-IDR2-DEP versus PDZ-pIDR2-DEP (**Fig. 4D, S7, Suppl. Tables 1 and 2**). Phosphorylation of IDR2 induced an important and significant decrease in the deuteration of the DEP domain. This protection is most likely due to a decreased solvent accessibility caused by the intramolecular contacts with pIDR2, as suggested by NMR. Slight protection was also observed for the PDZ domain (**Fig. 4Di**), but averaging all uptake plots resulted in no significant impact of the IDR2 phosphorylation on the deuterium uptake of the PDZ domain (**Fig. S7**).

Interestingly, the binding epitopes deduced by the two orthogonal methods, NMR and HDX-MS, overlap largely around a positively charged cleft of the DEP domain (**Fig. 4Eii**). Given that CK1ε multisite phosphorylation increases dramatically the negative charge density of the IDR2 proximally to DEP (**Fig. 4Ei**), the DVL3 conformational rearrangements should be charge-dependent. To test this, we analyzed the DVL3 conformation as a function of ionic strength by monitoring the chemical shifts in PDZ-pIDR2-DEP. At low salt (50 mM KCl in **Fig. 4Ai**, 0-100 mM NaCl in **Fig. S8**), DEP is coupled to pIDR2, but as the ionic strength increases, the DEP peaks move linearly until they coincide with those of PDZ-IDR2-DEP at high salt (**Fig. 4Aii, S8**). This is a clear indication that at high salt, DEP and pIDR2 are decoupled, revealing the pronounced electrostatic contribution to the intramolecular interaction (**Fig. 4Eii**).

Taken together, our data demonstrate that IDR2 phosphorylation couples order and disorder in the DVL3 continuum through electrostatically-driven contacts in a dynamic fashion, where the positively charged surface of DEP accommodates any of the adjacent phosphate groups. This implies that order-disorder coupling and conformational rearrangements in DVL3 depend on the negative charge density, charge clustering, or net charge proximally to DEP domain,^38^ features that are modulated by CK1ε-mediated phosphorylation in response to Wnt stimulus.

### Functional DVL3 phosphoswitch is required for the Wnt/β-catenin signal transduction

To better understand the role of the DEP proximal charge in DVL3 conformational switch, we examined mutants of the S/T cluster proximal to DEP by NMR. This phospho-switch mutant series included deletion of the cluster (DVL3 Δ S/T), phospho-preventive substitutions (DVL3 S/T-A), and phospho-mimetic substitutions (DVL3 S/T-E) of 12 S/T sites residing therein (**Fig. 5A**). Mutant DVL3 PDZ-IDR2-DEP proteins were produced in both non-phosphorylated and phosphorylated forms similarly to WT. The NMR data showed that, in all cases, the remnant S/T sites flanking the mutated proximal cluster (S378 and T398 in WT DVL3), as well as the distal cluster (T369 and T371), were phosphorylated by CK1ε (**Fig. S9**). Next, the spectroscopic behavior of the mutant proteins was analyzed using four peaks of the DEP domain that were well-resolved in the spectra (T459, T480, K483, S487) and experienced the largest CSPs in WT due to pIDR2 binding. Interestingly, the reporting peaks in mutants laid along the linear trajectories defined by the WT peaks in non-phosphorylated and phosphorylated forms, allowing to draw mechanistic conclusions on the DVL3 phospho-switch (**Fig. 5B**). Deletion of the proximal cluster in DVL3 (Δ S/T) shifted the distal cluster proximally to DEP and phosphorylation by CK1ε resulted in identical but weaker contacts with DEP, suggesting a partial conformational switch. This is attributed to the negative charge density of the phosphorylated S/Ts in the newly generated cluster proximally to DEP comprising only six S/T sites as opposed to the fourteen, densely spaced, S/T sites in WT. In DVL3 S/T-A, DEP remained uncoupled to IDR2 regardless of the phosphorylation, indicating that in a hydrophobic context proximal to DEP, phosphorylation of the distal cluster is not sufficient to couple DEP and IDR2. On the contrary, in DVL3 S/T-E, DEP and IDR2 were coupled *ab initio* due to the negative charge present proximally to DEP, but when the remaining S/T sites got phosphorylated, some additional rearrangements in the electrostatic contacts were observed. The experimentally determined behavior of the DVL3 phospho-switch mutant series is schematized in **Fig. 5Ci**.

**Figure 5.**
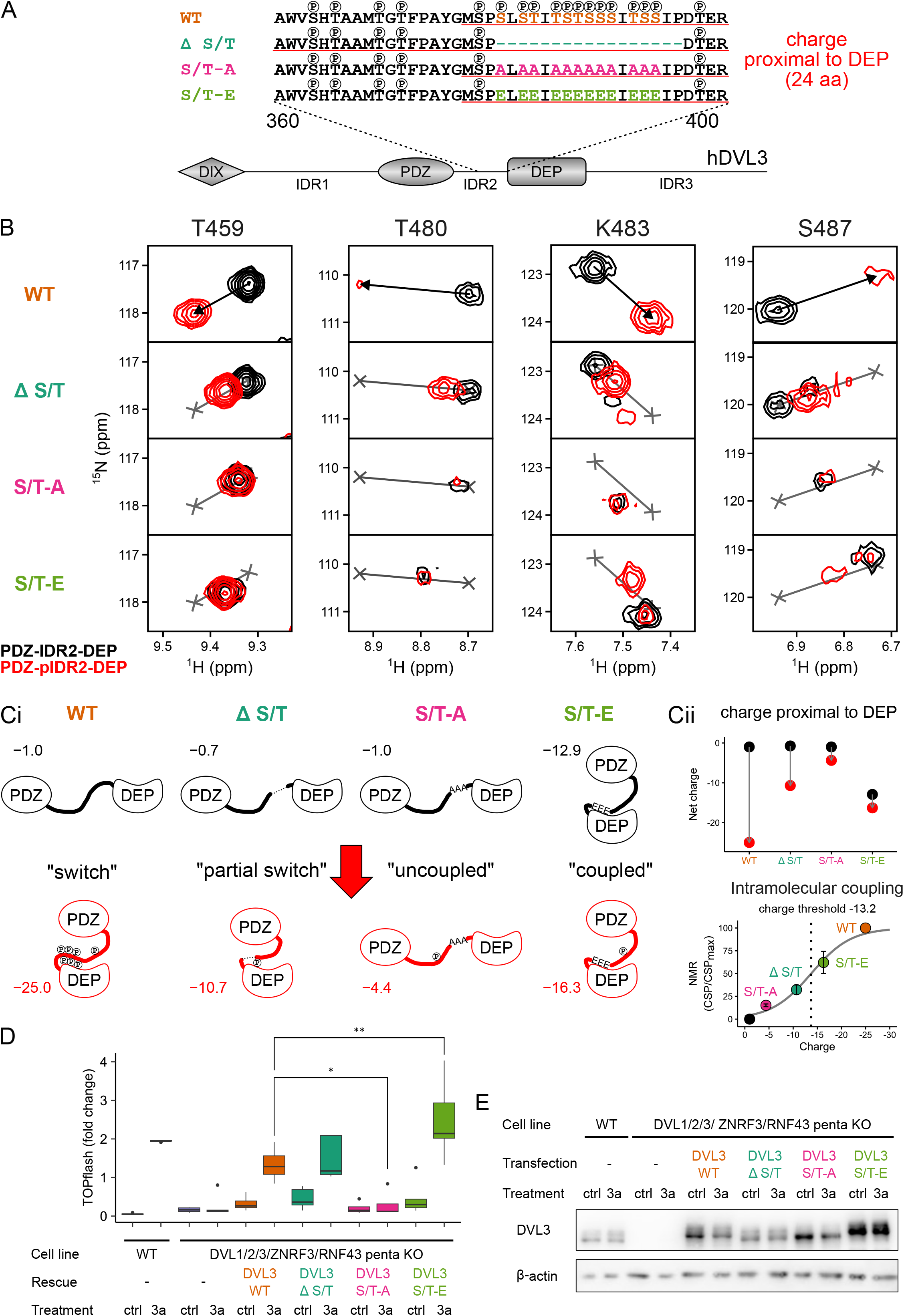
Charge of phospho-switch mutants affects intramolecular coupling and Wnt/β-catenin signal transduction. **A**. Sequences of WT and phospho-switch mutant series. Experimentally determined phosphorylation sites are marked on top of the S/T residues. The sequences used to calculate the net charge proximal to DEP are underlined in red. **B**. Overlay of ^1^H-^15^N HSQC spectra of hDVL3 PDZ-IDR2-DEP (black) and PDZ-pIDR2-DEP (red) for WT and phospho-switch mutants for the selected reporter peaks. Black arrows indicate chemical shift perturbation (CSP) caused by CK1ε phosphorylation in WT and are depicted as a grey line in the spectra of phospho-switch mutants. The reference WT positions (grey crosses) are used to determine the magnitude of coupling relative to WT. **Ci**. Schematic representation of DVL conformations for WT and phospho-switch mutants in the non-phosphorylated (black) and phosphorylated (red) forms. The net charge proximal to the DEP domain is reported for every conformation. **Cii**. The change in the net negative charge proximal to the DEP domain is depicted for WT and phospho-switch mutants (top). The magnitude of intramolecular coupling for WT and phospho-switch mutants as a function of net charge proximal to DEP was fitted using a sigmoidal function to determine the charge threshold (bottom). **D**. TOPflash rescue assay. Wild-type (WT) or *DVL1/2/3/ZNRF3/RNF43* penta KO cells were treated with control (ctrl) or Wnt-3a conditioned medium to induce signal transduction. Full length WT or phospho-switch mutants were reintroduced into the penta KO cells to rescue the signal. Statistical significance was tested by ANOVA. * stands for 0.05% significance level, ** stands for 0.01 significance level. N = 5. **E**. Expression of hDVL3 variants used in D. β-actin was used as the loading control.

The DVL3 conformations sampled in the non-phosphorylated forms and the ones elicited (or not) by CK1ε phosphorylation (**Fig. 5Ci**) suggested that both the net negative charge and the charge density in DEP proximity are the determinants of the phospho-switch at the molecular level (**Fig. 5Cii**). To better determine the switch-like behavior proposed by the data, we analyzed the intramolecular coupling between phosphorylated IDR2 and DEP with regard to the net negative charge in DEP proximity (**Fig. 5Cii, S10**). The output is a sigmoidal function of a multistep process describing the net charge accumulation in DEP proximity due to IDR2 phosphorylation, which is able to generate an ultrasensitive response in terms of the DEP conformational change. The model predicts a charge threshold of –13 at pH=6.5, which under physiological conditions (pH=7.2), would require 6-7 phosphorylations in the DEP proximal cluster to elicit the DVL3 conformational change (intramolecular coupling) and perhaps trigger signal transduction in response to Wnt stimulus.

To address whether there is a link between the functional phospho-switch and the capacity to transduce Wnt-3a signal in the Wnt/β-catenin pathway, we analyzed functionally the DVL3 phospho-switch mutant series using the TOPflash assay. Rescue experiments were performed using HEK293-T-REx *DVL1/DVL2/DVL3* triple KO cells^6^ as well as HEK293-T-REx penta KO cells (*DVL* triple KO, *RNF43* KO, *ZNRF3* KO)^39^, which lack all DVL paralogs in addition to E3 ligases RNF43 and ZNRF3, which attenuate the Wnt pathway at the receptor level (**Fig. 5D, S11A**). The results were nearly identical in both cell lines. *DVL* KO cells do not transduce Wnt-3a signal, a phenotype that is rescued by re-expression of exogenous DVL3 and quantified by TOPflash reporter assay.^5,6^ Intriguingly, DVL3 S/T-A mutant, in contrast to DVL3 WT that served as a positive control, completely failed to rescue TOPflash response to Wnt-3a. On the contrary, both DVL3 Δ S/T and S/T-E mutants were functional and could rescue the signal. However, none of these mutants could activate the pathway in the absence of Wnt-3a under the given experimental conditions. The level of expression of hDVL3 in lysates from the TOPflash experiment was similar, as analyzed by Western blotting (**Fig. 5E, S11B**). The rescue experiments support that charge accumulation proximal to the DEP domain and activation of the phospho-switch is a key functional event required, but not sufficient, for the Wnt-3a-induced signal transduction.

### DVL3 conformational phospho-switch controls the interaction with FZD receptors

To gain a better insight into the DVL3 phospho-switch role in Wnt signaling, we analyzed the DVL3 interactome by using a proximity-based proteomics approach. We created T-Rex-based cell lines expressing upon doxycycline induction WT or phospho-switch DVL3 mutants (**Fig. 6A**) fused to a promiscuous biotin ligase (TurboID)^40^ at protein levels close to endogenous DVL (**Fig. S12A**). Upon biotin supplementation, the TurboID biotinylates proteins in its proximity, which are, after cell lysis and streptavidin-based pulldown, subsequently identified by MS/MS (**Fig. 6A, Fig. S12A**).

**Figure 6.**
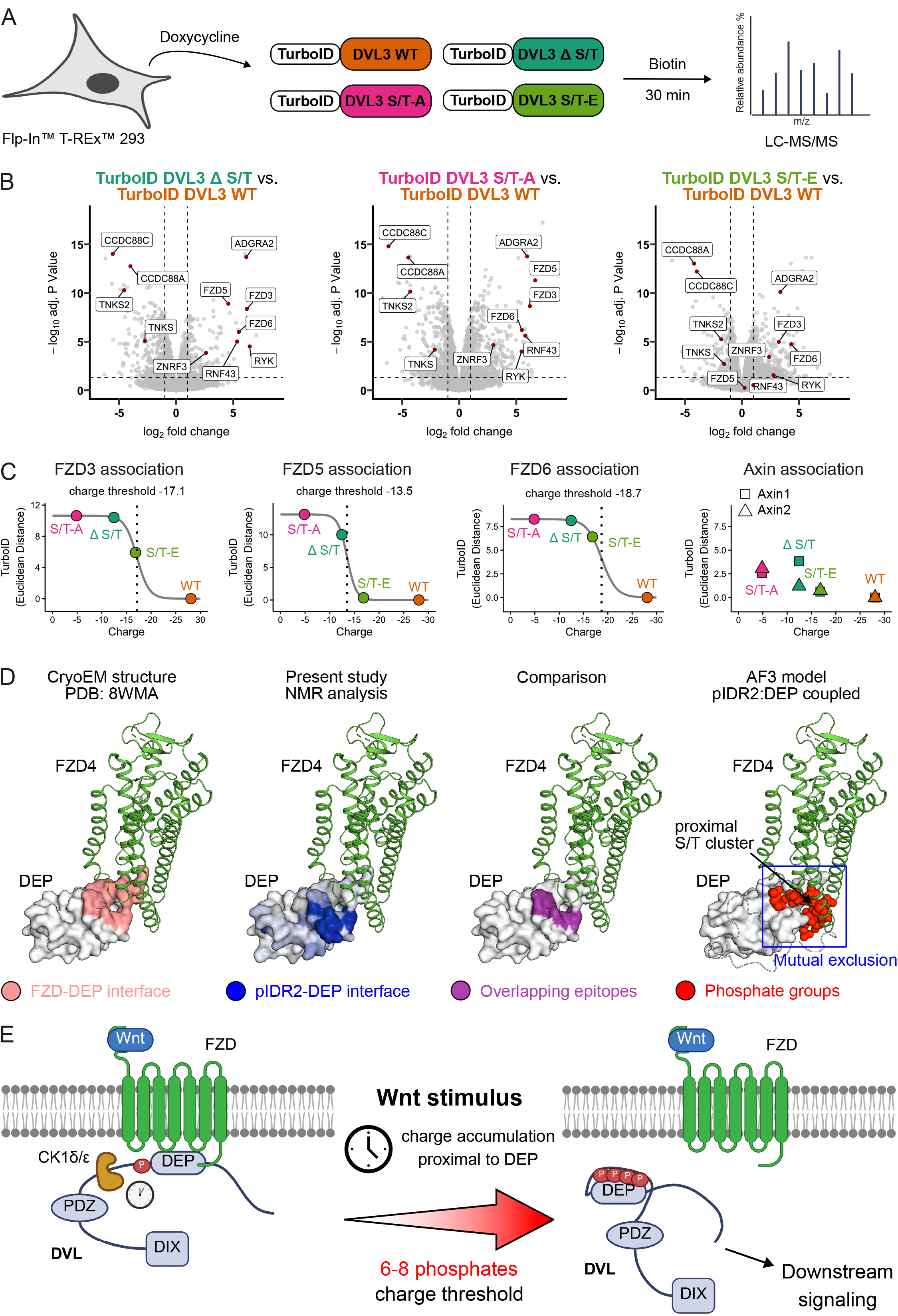
Phospho-switch regulates DVL interaction with FZDs. **A**. Schematic representation of the proximity-labelling experimental workflow. Expression of the phospho-switch mutant series fused to a promiscuous biotin ligase (TurboID) was induced by doxycycline. Biotinylated proteins were identified by MS/MS. **B**. Volcano plots comparing proteins enriched (right) or depleted (left) in the interactomes of phospho-switch mutants in comparison to the WT. **C**. The potency of each interaction was calculated as the Euclidean distance from the origin of the volcano plots comparing the interactome of phospho-switch mutants with WT. The corresponding values were plotted as a function of the net charge (at pH=7.2) proximal to the DEP domain and fitted using a sigmoidal function to determine the charge threshold for each FZD. Axin interaction with DVL3 did not show significant difference between the phospho-switch mutants and WT. **D**. FZD-DEP interface, pIDR2-DEP interface, and their overlap mapped onto the FZD-DEP structure (PDB ID: 8WMA). The pIDR2-DEP AlphaFold model superimposed onto the FZD-DEP structure (PDB ID: 8WMA). **E**. Mechanistic model of DVL phospho-switch in Wnt signaling. Wnt stimulus leads to the accumulation of negative charge proximal to the DEP domain. DVL intramolecular coupling competes with FZD binding, and when the charge threshold is reached, DVL dissociates from FZD, and transduces the Wnt signal to downstream components.

The interactome of WT and phospho-switch DVL3 mutants showed comparable interactions with the known and well-characterized DVL3 binding partners (**Fig. S12B**) such as other DVL paralogs (DVL1 and DVL2), AXIN1 and AXIN2, VANGL1 and VANGL2 as well as CK1δ (CSNK1D) and CK1ε (CSNK1E). This demonstrates that DVL3 phospho-switch mutants are not mislocalized or misfolded and interact with the same core subset of Wnt pathway components. In agreement, the immunocytochemistry experiment showed no differences on DVL3 subcellular localization in the presence or absence of exogenous CK1ε and their ability to colocalize with Axin in cytoplasmic puncta (**Fig. S11C**).

Interestingly, the unbiased comparison of DVL3 WT and phospho-switch series interactomes uncovered some remarkable differences (**Fig. 6B**). TurboID-DVL3 phospho-switch mutants biotinylated much more efficiently FZD proteins – FZD3, FZD5, and FZD6 – as well as other known transmembrane receptor complex proteins such as ADGRA2/GPR124,^41–43^ RYK^44,45^ or FZD-associated E3 ligases RNF43 and ZNRF3.^46–48^ On the other hand, the DVL3 variants with the phospho-switch mutations interacted less with several proteins associated with the downstream signaling – either in the Wnt/β-catenin pathway – such as TNKS and TNKS2,^49^ or in the non-canonical Wnt pathway – CCDC88A/Girdin and CCDC88C/Daple^50,51^. This demonstrates that the DVL3 mutants with fully or partially compromised phospho-switch interact with receptor complexes either with higher affinity or have a longer residence time on the receptors, which in turn attenuates their interaction with the downstream effectors.

Of particular interest to our analysis were the FZD receptors, since they engage physically with the DEP domain in recruiting DVL to the membrane after Wnt stimulus.^7,14^ The potency to interact with FZD receptors differed among individual phospho-switch DVL3 mutants. When the interaction with FZD receptors was analyzed as a function of the net negative charge in the phosphorylated region proximal to DEP, it correlated in the form of sigmoidal threshold curves (**Fig. 6C**). These traits align with the conformational coupling deduced by the analysis of the DVL3 phospho-switch series at the molecular level (**Fig. 5Cii**). The charge threshold (pH=7.2) reporting on FZD dissociation varies between FZDs but is in good agreement with the one determined by NMR data. Similar behavior was observed for other trans-membrane proteins associated with FZD receptor complex such as ADGRA2/GPR124, RYK, RNF43 and ZNRF3 (**Fig. S12C**). Importantly, other functionally relevant proteins (TNKS, TNKS2, CCDC88A, CCDC88C) followed opposite trend to FZDs, i.e. the higher the negative charge proximal to DEP, the better the binding to DVL3 (**Fig. S12C**), identifying these proteins as candidate effectors that recognize phosphorylated DVL after release from FZD. Interestingly, the majority of DVL3 interactors, including the absolutely essential downstream effectors AXIN1 and AXIN2, did not show a different binding to DVL3 phospho-switch variants (**Fig. 6C, Fig. S12C**). DVL3 phospho-switch thus controls only a subset of protein-protein interactions but does not affect other – especially those requiring other parts of DVL, such as the critical AXIN-DVL interaction that is mediated by DIX domain heteropolymerization.^9,18^

Altogether these data suggest a causal link between the intramolecular DVL coupling and loss of the interaction with the FZD receptors. Given the fact that the mutant with the completely deficient phospho-switch, DVL3 S/T-A, was incapable of Wnt-3a signal transduction, we conclude that IDR2 phosphorylation, followed by DVL3 conformational coupling is an essential step required for DVL to dissociate from FZDs and to signal towards downstream effectors. However, activation of the DVL phospho-switch and release from FZDs is not sufficient for pathway activation, which directly implicates that an additional step – such as the AXIN-mediated LRP5/6 recruitment and GSK3 inhibition in case of Wnt/β-catenin pathway, is required. A two-step mechanism is able to separate FZD activation from pathway output, and provides a general framework for signaling specificity within the Wnt pathway.

## Discussion

Here, we identified a Wnt-induced phosphorylation-dependent mechanism that is essential for DVL to act as the transducer in the Wnt pathway. CK1-mediated multisite phosphorylation of DVL IDR2 induces a charge-driven conformational change in DVL. DVL phosphorylation increases dramatically the negative charge density of IDR2 and promotes an allovalent electrostatic interaction with the positively charged cleft of the DEP domain. The DVL3 mutant (S/T-A), which is not amenable to switching conformations, is unable to transduce the Wnt signal in the Wnt/β-catenin pathway. Interestingly, the DVL3 mutant (S/T-E), in which IDR2 is coupled to DEP regardless of CK1 phosphorylation, did not behave as constitutively active but required a Wnt stimulus for activity. This suggests that the DVL conformational change driven by the intramolecular coupling between pIDR2 and DEP is required but not sufficient for downstream signal transduction.

The interactome analysis of the DVL3 mutant series suggested that the conformational phosphoswitch modulates the DVL interaction with the FZD receptor. A recent cryoEM structure has revealed that DEP engagement with FZD4 is mediated by two patches of interactions. (**Fig. 6D**).^11^ The first patch comprises the hydrophobic surface of the DEP finger that contacts the intracellular cavity of FZD4, and the second patch involves polar interactions with the positively charged cleft of DEP. When compared to the intramolecular coupling induced by DVL phosphorylation, the two DEP binding epitopes overlap, indicating that the two interactions are mutually exclusive, as supported by an AlphaFold 3 model of the pIDR2-DEP interaction. Therefore, we propose a mechanism in which Wnt-induced DVL conformational phospho-switch outcompetes FZD binding and, as an essential step required for the activation of the Wnt/β-catenin pathway, triggers DVL detachment from FZD (**Fig. 6E**).

Our data and analysis, may also offer the mechanistic link between FZD receptors and downstream signaling components. It has been proposed that the DEP domain of DVL undergoes a transition from monomers to domain-swapped dimers in relaying canonical Wnt signals.^7,52^ However, DEP domain swapping involves structural rearrangements around the DEP finger, and thus, it is incompatible with FZD binding.^11^ The DVL phospho-switch offers an elegant solution for the transition from DVL monomers bound to FZD to DVL swapped dimers, or larger oligomers, by detaching DEP from the receptor. Our data also provide both experimental evidence and a molecular mechanism for the hypothesis proposed by Ma and Kirschner^53^ based on the quantification of membrane-recruited DVL oligomers by TIRF microscopy. In their view, Wnt signal stabilizes larger oligomers of DVL, and as the DVL dwell time on the membrane increases, the DVL phosphorylation load may increase as well. At some point the phosphorylated DVL leaves the membrane to signal downstream. Multisite phosphorylation^13^ provides a solution that, in principle, displaces DVL from FZD only after the required phosphorylation threshold is reached. If the dwell time of DVL at the membrane – determined by the oligomerization status, which is directly correlated with Wnt concentration and the intensity of FZD clustering – is the key factor for activation,^53^ then the conformational phospho-switch described in this study, represents a perfect mechanism to define it. It is thus tempting to speculate that a combination of parameters, which define FZD clustering (Wnt-FZD affinity), pIDR2-DEP molecular coupling (e.g. number of S/T sites, their proximity to DEP and their density among the three DVL paralogs) and DVL-FZD interactions (DEP binding affinity to FZD paralogs), exert exquisite control over the downstream response. Our data provide a rationale for such scenario: TurboID interactomics (**Fig. 6C**) suggests different charge threshold for FZD5 that responds to Wnt-3a to trigger Wnt/β-catenin pathway,^54^ as compared to FZD3 and FZD6 that respond to Wnt-5a to trigger non-canonical Wnt pathway.^55,56^

At the molecular level, the phospho-switch is electrostatic in nature. The pIDR2 affinity for DEP is modulated by the number of phosphorylated S/T residues and their proximity to DEP. Proximity contributes to the binding by increasing the effective concentration through a favorable entropy.^57,58^ Yet, it is the cumulative electrostatic interactions that drive an ultrasensitive switch-like response when a threshold level of phosphorylation (and negative charge) is reached. To generate the maximum amount of ultrasensitivity, theory suggests that half of the sites are required to be phosphorylated for activation.^59^ Our NMR-based model predicted a charge of –13 for the response, corresponding to 7-8 phosphorylation events (at pH=6.5) amongst the fourteen S/T sites residing in the DEP proximal cluster. Mechanistically, an interesting implication of the cluster-based DVL multisite phosphorylation is that the response depends on the amount of phosphorylation (charge threshold), rather than the exact S/T sites being phosphorylated.

It remains to be determined how conserved the mechanism described in this study is. Sequence differences exist among IDR2 of DVLs across individual paralogs and species. For example, in the fruit fly *Drosophila melanogaster*, one of the most commonly used models of Wnt signaling, the dDSH IDR2 contains only 6 S/T residues – in comparison to hDVL3, which has 21 (**Fig. S13A**). However, our data still support the possibility that the same conformational switch is functional even in flies. More specifically, the sequence of dDSH IDR2 resembles that of IDR2 from hDVL3 Δ S/T and the charge profile of dDSH is similar to Δ S/T mutant (**Fig. S13B**), which showed a partial switch behavior and activity in the Wnt/β-catenin pathway. This suggests that dDSH contains the prototype version of the switch that has expanded in higher eukaryotes into a dense S/T cluster to allow for exquisite control over the signaling output.

In conclusion, we have identified a novel regulatory mechanism in Wnt signal transduction that links DVL phosphorylation to functionally significant conformational changes with direct consequences for the interaction with FZD. A mechanism that may be universal across DVL orthologs.

## Supporting information

All suplementary figures

## Acknowledgement

This project is supported by the Czech Science Foundation (GA23-06913S, GA22-25365S) and the Ministry of Education, Youth and Sports within programme Inter-Excellence II, INTER-ACTION (LUAUS25170). CIISB, Instruct-CZ Centre of Instruct-ERIC EU consortium, funded by MEYS CR infrastructure projects LM2023042 and CZ.02.01.01/00/23_015/0008175, is gratefully acknowledged for the financial support of the measurements at the CEITEC Josef Dadok National NMR Centre and Proteomics Facility. Computational resources were provided by the e-INFRA CZ project (ID:90254), supported by MEYS CR. Native MS and HDX-MS experiments were carried out using the facilities of the Montpellier Proteomics Platform (PPM, BioCampus Montpellier), supported by the regional funds FEDER/Région Occitanie, MUSE and the Labex EpiGenMed. The project National Institute for Cancer Research (Programme EXCELES, ID Project No. LX22NPO5102) - Funded by the European Union – Next Generation EU, is gratefully acknowledged for the financial support.

Alka Kumari Jadaun is gratefully acknowledged for her assistance in the preparation of S/T cluster mutant plasmids. Schematics of experimental procedures were created with BioRender.com.

## Author contributions

Conceptualization: M.M., J.K., S.B., K.T., V. Bryja; Data curation: K.H., C.B., H.P., K.G.; Formal analysis: M.M., J.K., P.P., K.H., C.B., H.P., O.Š., K.G., V. Bystrý; Funding acquisition: C.B., Z.Z., K.T., V. Bryja; Investigation: M.M., J.K., P.P., Z.H., K.H., C.B., O.Š., E.D.N. T. Č.; Methodology: M.M., K.H., M.K., Z.Z., K.T., V. Bryja; Project administration: Z.Z., K.T., V. Bryja; Computer scripts for data analysis: K.G.; Resources: T.G., S.B., C.B., M.K., Z.Z., K.T., V. Bryja; Supervision: K.T., V. Bryja; Visualization: M.M., P.P., C.B., O.Š.; Writing, review and editing: M.M., K.T., V. Bryja

## Declaration of interests

The authors declare no competing interests.

## Figure Legends

**Figure S1. Gels used for the analysis of phosphorylation of the endogenous hDVL3**. Black arrows represent hDVL3, while red arrows represent its phosphorylated form.

**Figure S2. Plots of all detected phosphorylation sites in endogenous hDVL3**. Different shapes represent different replicates. The mean values are indicated by black dashes. N = 3.

**Figure S3. *In vitro* multisite phosphorylation of hDVL3 IDR2 by CK1δ and CK1ε. A**. Progressive accumulation of multisite-phosphorylated species of hDVL3 IDR2 over the CK1ε-mediated phosphorylation time course as determined by MALDI-MS. Each sample was analyzed four times, and the mean value was plotted. **B**. LC-MS/MS detects the non-phosphorylated but not the phosphorylated peptides corresponding to the S/T cluster proximal to DEP. When the hDVL3 pIDR2 is dephosphorylated by protein phosphatase (PP) LC-MS/MS detects non-phosphorylated and phosphorylated peptides. MALDI-MS analysis of the same samples demonstrates that LC-MS/MS fails to detect highly phosphorylated peptides but is able to detect the corresponding peptides carrying 0, 1, or 2 phosphorylations after pIDR2 dephosphorylation. **C**. Overlay of ^1^H-^15^N HSQC spectra of native hDVL3 IDR2 (aa 335-396, black) and hDVL3 IDR2 phosphorylated *in vitro* by CK1δ (cyan). Phosphorylation is observed as a distinctive change in the chemical shift of S/T residues highlighted by black arrows. **D**. Correlation between CK1ε and CK1δ phosphorylation rates of individual IDR2 residues.

**Figure S4. Graphs representing phosphorylation time course of individual S/T residues of hDVL3 IDR2**. Phosphorylated by CK1ε (red) or CK1δ (cyan). Kinetics were measured as the decay of the peak intensity of the non-phosphorylated S/T residues.

**Figure S5. Graphs representing phosphorylation time course of residues neighboring S/T of hDVL3 IDR2**. Phosphorylated by CK1ε (red) or CK1δ (cyan). Kinetics were measured as the decay of the peak intensity of the non-phosphorylated residues.

**Figure S6. Analysis of IDR2 and pIDR2 interaction with the adjacent domains PDZ and DEP. A**. Overlay of ^1^H-^15^N HSQC spectra of ^15^N-labeled 50 µM hDVL3 PDZ domain (aa 243-351) free and in the presence of 250 µM hDVL3 IDR2 (aa 335-396). Overlay of ^1^H-^15^N HSQC spectra of ^15^N-labeled 50 µM hDVL3 IDR2 free and in the presence of 300 µM hDVL3 PDZ. CSP quantification against the PDZ or IDR2 sequence, respectively. **B**. Overlay of ^1^H-^15^N HSQC spectra of ^15^N-labeled 50 µM hDVL3 DEP domain free and in the presence of increasing amounts of hDVL3 IDR2. Overlay of ^1^H-^15^N HSQC spectra of ^15^N-labeled 50 µM hDVL3 IDR2 free and in the presence of increasing amounts of hDVL3 DEP. CSP quantification against the CSPs throughout DEP or IDR2, respectively. **C**. Overlay of ^1^H-^15^N HSQC spectra of ^15^N-labeled 50 µM hDVL3 PDZ domain free and in the presence of 300 µM phosphorylated hDVL3 IDR2 (pIDR2). Overlay of ^1^H-^15^N HSQC spectra of ^15^N-labeled 50 µM hDVL3 pIDR2 free and in the presence of 300 µM hDVL3 PDZ. CSP quantification against the PDZ or IDR2 sequence, respectively. **D**. CSPs in the DEP spectrum caused by the serial addition of phosphorylated IDR2 mapped onto the DEP structure (see also **Fig. 4B**).

**Figure S7. HDX-MS Coverage maps and heatmaps for the PDZ and DEP domains**. Blue bars represent the identified peptides after data curation, and the heatmaps represent the % differential relative fractional uptake (phosphorylated – non-phosphorylated PDZ-IDR2-DEP) for all deuteration time points analyzed (top to bottom: 0.5, 5, 10, 20, 40, 60, 120, and 240 min).

**Figure S8. Increasing ionic strength decouples DEP and pIDR2. A**. The DEP reporter peaks from the ^1^H-^15^N HSQC spectra of hDVL3 PDZ-IDR2-DEP (aa 243-496, black) overlaid with ^1^H-^15^N HSQC spectra of phosphorylated PDZ-IDR2-DEP (PDZ-pIDR2-DEP, red) in the presence of increasing NaCl concentrations. Arrows indicate the chemical shift perturbations (CSPs) of the selected residues caused by phosphorylation at different salt concentrations. **B**. CSPs of the selected residues shown in **A** as a function of increasing NaCl concentration. The following concentrations were tested: 0 mM, 50 mM, 100 mM, 250 mM, and 500 mM NaCl.

**Figure S9. NMR analysis of WT and phospho-switch mutant series**. Overlay of ^1^H-^15^N HSQC spectra of PDZ-IDR2-DEP and PDZ-pIDR2-DEP. DEP reporter peaks are circled and indicated by arrows. Zoom-in view highlights the G370 chemical shift change reporting on T369-T371 phosphorylation and annotated the peak position of the phosphorylated T398 (pT398). Phosphorylation-induced CSPs are plotted against the DVL3 sequence. **A**. WT. **B**. Δ S/T. **C**. S/T-A. **D**. S/T-E.

**Figure S10. Intramolecular coupling measured by NMR as a function of net charge in the proximity of the DEP domain. A**. Plots for individual reporter peaks T459, T480, K483, and S487, respectively, reporting on intramolecular coupling as a function of net charge proximal to DEP fitted to a sigmoidal function. **B**. Overlay of ^1^H-^15^N HSQC spectra of S/T-A in non-phosphorylated and phosphorylated form and the DEP domain of WT DVL3. S/T-A peaks K483 and S487 report on the exchange phenomena observed in DEP domain which are unrelated to IDR2 phosphorylation and intramolecular coupling. Thus, the S/T-A data points for K483 and S487 indicated by arrows in **A** were not used in calculating the mean value of coupling in the corresponding plot in **Fig. 5Cii**.

**Figure S11. Cellular effects of the phospho-switch mutant series. A**. TOPflash assay. Wild-type (WT) or *DVL1/2/3* triple KO cells were treated with control (ctrl) or Wnt-3a conditioned media to induce signal transduction. Full length WT or phospho-switch mutant series were reintroduced into the triple KO cells to rescue the signal. Statistical significance was tested by ANOVA. ** stands for 0.01 significance level. N = 4. **B**. Expression of hDVL3 variants used in A. β-actin was used as the loading control. **Ci**. Representative pictures of the subcellular localization of hDVL3 mutant variants, overexpressed alone or together with CK1ε or Axin1, respectively. Scale bar = 10 µm **Cii**. Quantification of the subcellular localization of hDVL3 mutant variants overexpressed alone or together with CK1ε, respectively. N = 3. **Ciii**. Quantification of the colocalization of hDVL3 mutant variants with Axin1. N = 1.

**Figure S12. TurboID validation. A**. Expression of TurboID hDVL3 variants induced by 1 µg/ml doxycycline, and TurboID biotinylation induced by 50 µM biotin, analyzed by western blotting. β-actin was used as the loading control. **B**. Volcano plots showing specific interactions for WT and phospho-switch DVL3 mutants. Volcano plots show comparison between TurboID-DVL3 and TurboID only. Selected well know DVL3 interactors acting in the Wnt pathway are highlighted. **C**. The potency of selected proteins to interact with WT DVL3 and phospho-switch variants was calculated as the Euclidean distance from the origin of the volcano plots comparing the interactome of phospho-switch mutants with WT. The corresponding values were plotted as a function of the pIDR2 net charge (at pH=7.2) proximal to the DEP domain.

**Figure S13. Δ S/T and dDSH similarities. A**. Sequence alignment of IDR2 of human DVL3 (WT and Δ S/T mutant) and *Drosophila melanogaster* dDSH. Conserved S/T residues are highlighted in dark grey and other negatively charged residues in dDSH are shown in bold. **B**. Local charge distributions of PDZ-IDR2-DEP and PDZ-pIDR2-DEP for hDVL3 Δ S/T and dDSH.

## EXPERIMENTAL MODEL

### Human Cell lines

FreeStyle™ 293-F (RRID:CVCL_D603) cells were cultured at 37 °C, while shaking in a plastic Erlenmeyer flask, in FreeStyle™ 293 Expression Medium (12338002, Thermo Fisher Scientific) supplemented with 1% penicillin-streptomycin (XC-A4122/100, Biosera). To stimulate the Wnt pathway and analyse the phosphorylation status of endogenous DVL3, two hundred million (200 x 10^6^) cells in a total volume of 100 ml were cultivated overnight with 1 µM Porcupine inhibitor (LGK974, MedChemExpress) one day prior to the experiment to block autocrine Wnt stimulation. The next day, cells were treated with 50 % of the volume Wnt-3a or Wnt-5a or control conditioned media for 2 hours together with 10 % of the volume RSPO conditioned media. After the treatment, cells were centrifuged at 400 g for 5 min at 4 °C. The cell pellet was shock frozen with liquid nitrogen and stored for future use at -80 °C.

HEK T-REx-293 cells (RRID:CVCL_D585), HEK T-REx-293 DVL triple KO^6^ and HEK T-REx-293 penta KO (DVL triple KO, RNF43 KO, ZNRF3 KO)^39^ cells and TurboID cell lines were cultured at 37 °C in Dulbecco’s modified Eagle’s medium (DMEM; 41966–029, Gibco, Life Technologies) supplemented with 10% fetal bovine serum (FBS; 10270–106, Gibco, Life Technologies), 1% penicillin-streptomycin (XC-A4122/100, Biosera), and 1% L-glutamine (25030024, Life Technologies). HEK T-REx-293 DVL triple KO and penta KO cells were generated from HEK T-REx-293 cells using CRISPR-Cas9.

TurboID stable cell lines expressing either WT DVL3 or the phospho-switch mutant series N-terminally fused with TurboID, and a control cell line expressing TurboID alone were generated from Flp-In™ T-REx™ 293 cell line (RRID:CVCL_U421) under Hygromycin B selection (ant-hg-1, Invivogen).

### Bacterial strains

*E. coli* BL21 (DE3) was used for protein expression and *E. coli* DH5α was used for plasmid propagation.

## METHOD DETAILS

### Conservation analysis of hDVL3

Conservation analysis was performed as described previously ^12^. Briefly, 854 sequences from 432 organisms were aligned using the MUSCLE algorithm^61^ of the msa package in R.

### Purification of endogenous hDVL3

The cell pellet was resuspended in 30 ml of binding buffer (containing 150 mM NaCl, 25 mM Tris [pH 8.0], 10 mM imidazole) with addition of 1% NP-40 (74385, Sigma-Aldrich) and protease inhibitor cocktail (11836145001, Roche) and sonicated for 2 minutes (using a pulse of 5 seconds on and 10 seconds off) at 4 °C. The cell lysate was centrifuged at 27,000 g for 1 hour at 4 °C. The supernatant was loaded onto 500 µl of nickel Sepharose beads (Ni Sepharose^TM^ 6 Fast FLow, GE Healthcare) equilibrated with the binding buffer. Beads were washed with the binding buffer; the protein was eluted with the biding buffer supplemented with 400 mM imidazole and 2 mM ethylenediaminetetraacetic acid (EDTA) was added to the sample. The eluate was incubated with 10 µg of DVL 3 antibody (sc-8027, Santa Cruz) for 1 hour at 4 °C, then overnight with Protein G Dynabeads™ (10004D, Thermo Fisher Scientific). Next day, the beads were washed with a buffer containing 150 mM NaCl, 20 mM Tris [pH 8.0] and 2 mM EDTA. The sample was resolved on an 8% polyacrylamide gel and stained with Coomassie brilliant blue (CBB).

### CK1 inhibition

10 µM MU1742^62^ was used to inhibit kinase activity of all CK1 isoforms.

### Molecular cloning

hDVL3 segments of interest (IDR2, aa 335-396; DEP, aa 398-496; PDZ, aa 243-351, and PDZ-IDR2-DEP, aa 243-496) were amplified by PCR and inserted into a pETM11 or pET-Z (IDR2) expression vectors via restriction digestion between the 5’-NcoI and 3’-KpnI sites, using BspHI (R0517, NEB) and KpnI (R3142S, NEB) restriction enzymes. The DNA segment encoding Casein kinase 1δ or ε was amplified by PCR and inserted into pETM11 expression vector via restriction digestion between the 5’-NcoI and 3’-EcoRI sites, using NcoI (R0193L, NEB) and EcoRI (R3101S, NEB) restriction enzymes. The DNA segment encoding Casein kinase 1ε core was excised from our in-house pETM11-CK1ε^63^ via restriction digestion using NcoI and EcoRI enzymes and subcloned into a pCDF-11 vector between the 5’-NcoI and 3’-EcoRI sites. Vectors were pretreated with Antarctic phosphatase (M0289, NEB), and the DVL, CK1ε, or CK1δ constructs were ligated using T4 DNA Ligase (M0202, NEB). The pETM11 and pET-Z vectors contain a kanamycin resistance marker, and the pCDF-11 vector contains a spectinomycin resistance marker. The pETM11 vector features an N-terminal His6-tag, followed by a TEV protease cleavage site before an inserted fragment. The pET-Z vector features an additional Z-tag inserted between the His6-tag and the TEV protease cleavage site before an inserted fragment. The pCDF vector does not contain any affinity chromatography tag. The sequences of the cloned inserts were verified through Sanger sequencing (Eurofins Genomics).

### Recombinant protein expression and purification

Chemically competent *E. coli* BL21 (DE3) cells, suitable for gene expression, were transformed with the DVL constructs in pET-Z vectors or pETM11 using a heat shock method. Bacteria were cultivated in either LB broth or M9 minimal medium. Minimal medium contained ^15^NH_4_Cl (CN80P10, Cortecnet) at 0.5 g/l as the sole nitrogen source for uniformly ^15^N-labelled proteins, or 0.5 g/l of ^14^NH_4_Cl (254134, Sigma-Aldrich) supplemented either with 200 mg/l ^15^N L-serine (NLM-2036-1, Cambridge Isotope Laboratories, Inc.) or 200 mg/l ^15^N/^13^C L-threonine (CCN3900P01, Cortecnet) together with 200 mg/l 2-ketobutyrate (K401, Sigma-Aldrich) and 400 mg/l glycine (14570-30500, Penta Chemicals Unlimited) for the ^15^N serine selective and ^15^N/^13^C threonine selective labelling, respectively. To obtain DVL phosphorylated proteins (pDVL), 200 ng of DVL pET-Z or pETM11 vectors were co-transformed with 600 ng of CK1ε pCDF vector. Bacteria were grown under an antibiotic selection (30 µg/ml of kanamycin and 100 µg/ml of spectinomycin for co-expression with CK1ε).

The cells were cultivated at 37 °C until OD_600_ reached around 0.8, then the temperature was lowered to 16 °C and the cells were allowed to adapt for one hour. Gene expression was induced by adding 0.25 or 0.5 mM of isopropyl β-D-1-thiogalactopyranoside (IPTG; A1008, PanReac AppliChem). After 16-30 hours of cultivation at 16 °C, cells were harvested by centrifugation at 4,000 g for 10 minutes at 10 °C. In the case of the PDZ, DEP, or PDZ-IDR2-DEP, pellets were resuspended in ice-cold lysis buffer (25 mM Tris [pH 8.0], 500 mM NaCl, 10 mM imidazole, 10% glycerol, 1% NP-40, and 300 mM sucrose). In the case of the CK1δ or ε, pellets were resuspended in ice-cold lysis buffer (25 mM Tris [pH 7.5], 500 Mm NaCl, 10 mM imidazole, 10% glycerol, 1% NP-40, and 300 mM sucrose). In the case of IDR2 peptide, pellets were resuspended in ice-cold urea buffer (6 M urea, 25 mM Tris [pH 8.0], 500 mM NaCl, 10 mM imidazole). Pellets were frozen at -20 °C for future use.

Cell pellets were thawed and supplemented with 2 mM β-mercaptoethanol (BME; 63689, Sigma-Aldrich), protease inhibitor cocktail, and phenylmethane sulfonyl fluoride (PMSF; 329-98-6, AppliChem). The cells were sonicated for 20 minutes (using a pulse of 5 seconds on and 10 seconds off) at 4 °C. The cell lysate was centrifuged at 27,000 g for 1 hour at 4 °C.

For the purification of phosphorylated or non-phosphorylated hDVL3 IDR2 peptides (aa 335-396), the supernatant was applied to the Ni-NTA column (HiTrap™ IMAC HP 5 ml, Cytiva), pre-equilibrated with the urea buffer. Subsequently, the urea buffer was exchanged for the binding buffer (25 mM Tris [pH 8.0], 500 mM NaCl, 10 mM imidazole) using gradient between the urea and the binding buffer. The peptide was eluted by a gradient between the binding buffer and the high imidazole buffer (25 mM Tris [pH 8.0], 500 mM NaCl, 500 mM imidazole). The His6-tag and Z-tag were cleaved by TEV protease at 4 °C overnight. Non-phosphorylated IDR2 peptide was dialyzed to the binding buffer, while phosphorylated IDR2 peptide was dialyzed to the buffer containing 50 mM NaCl, 25 mM Tris [pH 8.0], 5 % glycerol. Non-phosphorylated IDR2 peptide without the tag was collected from the flow-through of the second Ni-NTA step. Phosphorylated IDR2 peptide without the tag was applied to the Ni-NTA column coupled with the anion exchange. The peptide was eluted from the anion exchange column using a salt gradient from 50 mM to 1 M NaCl in a buffer of 25 mM Tris [pH 8.0] containing 1 mM EDTA. IDR2 peptides were then concentrated using ultrafiltration (Vivaspin 20, 5,000 MWCO PES, Sartorius) at 3,220 g at 10°C. In the final purification step, the IDR2 peptides were loaded into a Superdex 75 Increase 10/300 GL column (Cytiva) for size exclusion chromatography and eluted in 50 mM KCl, 50 mM phosphate buffer [pH 6.5].

For the purification of hDVL3 PDZ domain (aa 243-351)^12^ or DEP domain (aa 398-496) proteins, the supernatant was applied to a Ni-NTA column, pre-equilibrated with a binding buffer (25 mM Tris [pH 8.0], 500 mM NaCl, 10 mM imidazole). The protein was eluted by a gradient between the binding buffer and a high imidazole buffer (25 mM Tris [pH 8.0], 500 mM NaCl, 500 mM imidazole). The His6-tag was cleaved by TEV protease (purified in-house) at 4 °C overnight, and the sample was dialyzed to remove the imidazole. The sample was applied to the Ni-NTA column, now without any tag; the protein was collected in the flow-through of the second Ni-NTA step. The sample was concentrated using ultrafiltration (Vivaspin 20 5,000 MWCO PES, Sartorius) at 3,220 g at 22°C. In the final purification step, the sample was loaded into a Superdex 75 Increase 10/300 GL column for size exclusion chromatography and eluted in 50 mM KCl, 50 mM phosphate buffer [pH 6.5].

For the purification of phosphorylated or non-phosphorylated hDVL3 PDZ-IDR2-DEP protein (aa 243-496), WT and phospho-switch mutant series, the supernatant was applied to a Ni-NTA column, pre-equilibrated with a binding buffer (25 mM Tris [pH 8.0], 500 mM NaCl, 10 mM imidazole). The protein was eluted by a gradient between the binding buffer and a high imidazole buffer (25 mM Tris [pH 8.0], 500 mM NaCl, 500 mM imidazole). The His6-tag was cleaved by TEV protease (purified in-house) at 4 °C overnight, and the sample was dialyzed to remove the imidazole. The sample was applied to the Ni-NTA column, now without any tag; the protein was collected in the flow-through of the second Ni-NTA step. Subsequently, the protein was dialyzed to drop the salt concentration to 50 mM NaCl and applied to an anion exchange column (HiTrap™ Q HP 5 ml, Cytiva). The protein was eluted using a salt gradient from 50 mM to 1 M NaCl in a buffer of 25 mM Tris [pH 8.0] containing 1 mM EDTA and concentrated using ultrafiltration (Vivaspin 6, 10,000 MWCO PES, Sartorius) at 3,220 g at 10°C. The final purification step was size exclusion chromatography (Superdex 75 Increase 10/300 GL column), and PDZ-IDR2-DEP protein was eluted in 50 mM KCl, 50 mM phosphate buffer [pH 6.5]. The protein concentration was determined by absorbance at 280 nm using the appropriate extinction coefficient.

For the purification of CK1 isoforms, the supernatant was applied to a Ni-NTA column, pre-equilibrated with a binding buffer (25 mM Tris [pH 7.5], 500 mM NaCl, 10 mM imidazole). The protein was eluted by a gradient between the binding buffer and a high imidazole buffer (25 mM Tris [pH 7.5], 500 mM NaCl, 500 mM imidazole). The His6-tag was cleaved by TEV protease (purified in-house) at 4 °C overnight, and the sample was dialyzed to remove the imidazole and to drop the salt concentration to 50 mM NaCl, then applied to cation exchange column (HiTrap™ SP 5 ml, Cytiva). The protein was eluted using a salt gradient from 50 mM to 1 M NaCl in a buffer of 25 mM Tris [pH 7.5] containing 1 mM EDTA and concentrated using ultrafiltration (Vivaspin 6, 10,000 MWCO PES, Sartorius) at 3,220 g at 4°C. The final purification step was size exclusion chromatography (Superdex 75 Increase 10/300 GL column) the protein was eluted in 50 mM KCl, 50 mM phosphate buffer [pH 6.5]. The protein concentration was determined by absorbance at 280 nm using the appropriate extinction coefficient.

### LC-MS/MS analysis

Samples were loaded onto SDS-PAGE gels and the corresponding bands were excised. After destaining, the proteins in gel pieces were incubated with 10 mM Dithiothreitol (DTT) at 56 °C for 45 min. After the removal of excess DTT, samples were incubated with 55 mM iodoacetamide at room temperature in darkness for 30 min, then alkylation solution was removed and gel pieces were hydrated for 45 min at 4 °C in digestion solution (5 ng/µl trypsin, sequencing grade, Promega, in 25 mM ammonium bicarbonate). The trypsin digestion proceeded for 2 hours at 37 °C on Thermomixer (750 rpm; Eppendorf). Subsequently the tryptic digests were cleaved by chymotrypsin (5 ng/µl, sequencing grade, Roche, in 25 mM ammonium bicarbonate) for 2 hours at 37 °C. Digested peptides were extracted from gels using 50% ACN solution with 2.5% formic acid and concentrated in speedVac concentrator (Eppendorf). 1/10 or 1/2 (in the case of the endogenous hDVL3) of concentrated sample was transferred to LC-MS vial with pre-added polyethylene glycol (PEG; final concentration 0.001%)^64^ and directly analyzed by LC-MS/MS for protein identification

The rest of sample was used for phosphopeptide analysis. Phosphopeptides were enriched using High-Select™ TiO2 Phosphopeptide Enrichment Kit (A32993, Thermo Fisher Scientific) according to the manufacturer’s protocol and extracted into LC-MS vial with pre-added PEG (final concentration 0.001%). Resulting peptides were analysed by LC-MS/MS.

For the LC-MS/MS analysis of the PDZ-pIDR2-DEP phosphorylation, all peptide mixtures (with and without phosphoenrichment step) were analyzed using using Ultimate 3000 RSLCnano system connected to Orbitrap Elite hybrid spectrometer (Thermo Fisher Scientific) with ABIRD (Active Background Ion Reduction Device; ESI Source Solutions) and Digital PicoView 550 (New Objective) ion source (tip rinsing by 50% acetonitrile) installed. Prior to LC separation, digests were online concentrated and desalted using trapping column (300 μm × 5 mm, μPrecolumn, 5μm particles, Acclaim PepMap100 C18, heated compartment temperature 40 °C, Thermo Fisher Scientific). After washing of the trapping column with 0.1% FA, the peptides were eluted (flow 4 μl/min) from the trapping column onto Acclaim PepMap RSLC C18 column (3 µm particles, 75 μm × 500 mm; Thermo Fisher Scientific) by 65-min gradient. Mobile phase A (0.1% FA in water) and mobile phase B (0.1% FA in 80% acetonitrile) were used in both cases. The gradient elution started at 1% of mobile phase B and increased from 1% to 35% during the first 45 min, then increased linearly to 80% of mobile phase B in the next 10 min and remained at this state for the next 10 min. Equilibration of the trapping column and the analytical column was done prior to sample injection to sample loop.

MS data were acquired in a data-dependent strategy using a survey scan (350-2000 m/z). The resolution of the survey scan was 60 000 (400 m/z) with a target value of 1×106 ions, one microscan and maximum injection time of 1,000 ms. High resolution (resolution 15,000 at 400 m/z) HCD MS/MS spectra were acquired with a target value of 5×105. Normalized collision energy was 32% for HCD spectra. The maximum injection time for MS/MS was 500 ms. Dynamic exclusion was enabled for 45 s after one MS/MS spectra acquisition and early expiration was disabled. The isolation window for MS/MS fragmentation was set to 2.0 m/z.

LC-MS/MS analyses of all peptide mixtures (with and without phosphoenrichment step) of the endogenous DVL3 were done using Ultimate 3000 RSLCnano system connected to Orbitrap Exploris 480 spectrometer (Thermo Fisher Scientific) with EASY Spray ion source (Thermo Fisher Scientific) installed. Prior to LC separation, tryptic digests were online concentrated and desalted using trapping column (300 μm × 5 mm, μPrecolumn, 5μm particles, Acclaim PepMap100 C18, heated compartment temperature 40 °C, Thermo Fisher Scientific). After washing of the trapping column with 0.1% FA, the peptides were eluted (flow 6 μl/min) from the trapping column onto Acclaim PepMap RSLC C18 column (2 µm particles, 75 μm × 250 mm; Thermo Fisher Scientific) by 74-min gradient. Mobile phase A (0.1% FA in water) and mobile phase B (0.1% FA in 80% acetonitrile) were used in both cases. The gradient elution started at 3% of mobile phase B and increased from 3% to 37% during the first 64 min, then increased linearly to 80% of mobile phase B in the next 8 min and remained at this state for the next 2 min. Equilibration of the trapping column and the analytical column was done prior to sample injection to sample loop.

MS data were acquired in a data-dependent strategy using a survey scan (350-2000 m/z). The resolution of the survey scan was 120,000 (200 m/z) with a target value of 2.5×106 ions, one microscan and maximum injection time of 500 ms. High resolution (resolution 30,000 at 200 m/z) HCD MS/MS spectra were acquired with a target value of 2×105. Normalized collision energy was 30% for HCD spectra. The maximum injection time for MS/MS was 250 ms. Dynamic exclusion was enabled for 45 s after one MS/MS spectra acquisition and early expiration was disabled. The isolation window for MS/MS fragmentation was set to 1.2 m/z.

The analysis of the mass spectrometric RAW data files was carried out using the Proteome Discoverer software (Thermo Fisher Scientific; version 1.4) with the in-house Mascot (Matrixscience) search engine utilization. MS/MS ion searches were done against an in-house database containing expected protein of interest with additional sequences from cRAP database (downloaded from http://www.thegpm.org/crap/). Mass tolerance for peptides and MS/MS fragments was 7 ppm and 0.03 Da, respectively. Oxidation of methionine, deamidation (N, Q), phosphorylation (S, T, Y) and carbamidomethylation of C as optional modifications were set for all searches. The phosphoRS feature was used for phosphorylation localization. Peptides with false discovery rate (FDR; q-value) < 1%, rank 1 and with at least 6 amino acids were considered.

Quantitative information was assessed and manually validated in Skyline software (Skyline daily). Site occupancies were calculated as published in ^29^.

Bead bound protein complexes (in the case of the TurboID experiment) were processed and digested as described elsewhere.^65,66^ Digested peptides were evaporated completely in SpeedVac concentrator (Thermo Fisher Scientific). The resulting peptides were analyzed by LC-MS/MS.

LC-MS/MS analyses were done using Ultimate 3000 RSLCnano system connected to Orbitrap Exploris 480 spectrometer (Thermo Fisher Scientific) with EASY Spray ion source (Thermo Fisher Scientific) installed. Prior to LC separation, tryptic digests were online concentrated and desalted using trapping column (300 μm × 5 mm, μPrecolumn, 5μm particles, Acclaim PepMap Neo C18, Thermo Fisher Scientific). After washing of the trapping column with 0.1% FA, the peptides were eluted (flow 300 nl/min) from the trapping column onto Aurora C18 (75 μm × 250 mm, 1.7 μm particles, heated to 50°C, Ion Opticks) by 98-min gradient (mobile phase A: 0.1% FA in water; mobile phase B: 0.1% FA in 80% acetonitrile).

Data were acquired in a data-independent acquisition mode (DIA). The survey scan covered m/z range of 350-1,400 at a resolution of 60,000 (at m/z 200) and a maximum injection time of 55 ms (normalized AGC target 300%). HCD MS/MS (27% relative fragmentation energy) were acquired in the range of m/z 200-2,000 at 30,000 resolution (maximum injection time 55 ms, normalized AGC target 1000%). An overlapping window scheme in the precursor m/z range from 400 to 800 were used as isolation window placements. Raw data were converted to mzML format using msconvert (version 3.0.21193-ccb3e0136) using peakPicking (vendor msLevel=1-) and demultiplex (optimization=overlap_only massError=10ppm) filters applied.

DIA data were processed in DIA-NN^67^ (version 1.8.1) against iRT database (11 sequences in total) and UniProtKB Human (number of protein sequences: 20,590). No optional modifications, carbamidomethylation as fixed modification, and trypsin/P enzyme with 1 allowed missed cleavage and peptide length 7-30 were set during the library preparation. False discovery rate (FDR) control was set to 1% FDR. MS1 and MS2 accuracies as well as scan window parameters were set based on the initial test searches (median value from all samples ascertained parameter values). MBR was switched on.

Reported protein intensities were further processed using the software container environment (https://github.com/OmicsWorkflows). Processing workflow described briefly, it covered: a) removal of low-quality precursors and contaminant protein groups, b) precursor intensities normalization by loessF algorithm, c) precursor intensities imputation by global quantile (0.001), d) protein group MaxLFQ intensities calculation using iq R package^68^ (version 1.9.12) and log2 transformation, and e) differential expression analysis using LIMMA statistical test.

### MALDI-MS analysis

Samples in a volume of 0.6 µl were mixed with 2.4 µl of the MALDI matrix solution (12.5 mg/ml ferulic acid in water:acetonitrile:formic acid 50:33:17 v/v mixture), 0.6 µl of the mixture was spotted onto a stainless steel sample plate, and allowed to dry at room temperature.

MALDI-TOF mass spectra measurements were carried out using an ultrafleXtreme instrument (Bruker Daltonics, Bremen, Germany) operated in linear positive mode under FlexControl 3.4 software (Bruker Daltonics). External calibration of the mass spectra was performed using *E. coli* DH5 alpha standard (Bruker Daltonics). Four independent mass spectra, each comprising 1,000 laser shots, were acquired from each sample. Mass spectra were processed using Flex Analysis (version 3.4, Bruker Daltonics). The peak areas of individual protein forms were used to estimate their relative abundance.

### Native ESI-MS measurements

Prior to MS analysis, proteins were buffer exchanged into 200 mM ammonium acetate buffer [pH 7.4] (Sigma) using Bio-Spin microcentrifuge columns (Bio-Rad Laboratories). Intact mass spectra were recorded on a Synapt G2-Si HDMS instrument (Waters Corporation) modified for high mass analysis and operated in ToF mode. Samples were introduced into the ion source using borosilicate emitters (Thermo Scientific). Optimized instrument parameters were as follows: capillary voltage 1.4 kV, sampling cone voltage 80 V, offset voltage 50 V, collision voltage 50 V, transfer collision voltage 25 V, and argon flow rate 4 ml/min. Data was processed using MassLynx v.4.2 (Waters).

### *In vitro* phosphorylation and dephosphorylation

For the LC-MS/MS analysis of hDVL3 PDZ-IDR2-DEP, 5 µM of hDVL3 PDZ-IDR2-DEP (aa 243-496) was mixed with 125 nM of CK1ε (aa 1-301) in the buffer containing 200 mM NaCl, 50 mM Tris [pH 7.5], 1mM EDTA, 1 mM ATP, 10 mM MgCl_2_. For the MALDI-MS analysis, 50 µM of ^15^N-labelled hDVL3 IDR2 was mixed with 1 µM CK1ε (aa 1-301) in the buffer containing 50 mM KCl, 50 mM phosphate buffer [pH 6.5], 1mM EDTA and 2 mM ATP and 10 mM MgCl_2_. Both reactions took place at 25 °C.

For the dephosphorylation reaction, the protein was treated with 10 U of alkaline phosphatase (04571363103, Roche) for 2 hours at 37 °C in the buffer provided by the supplier.

### NMR spectroscopy

NMR ^1^H-^15^N HSQC spectra were measured on 700, 850 or 950 MHz Bruker Avance *NEO* spectrometers equipped with ^1^H/^13^C/^15^N TCI cryogenic probe heads with z-axis gradients. We measured 50 µM of ^15^N-labelled hDVL3 IDR2 (aa 335-396), phosphorylated hDVL3 IDR2, hDVL3 DEP (aa 398-496), hDVL3 PDZ (aa 243-351) or 90 µM of ^15^N-labelled DVL3 of non-phosphorylated or phosphorylated PDZ-IDR2-DEP (aa 243-496), WT and phospho-switch mutant series, in the buffer containing 50 mM KCl, 50 mM phosphate buffer [pH 6.5], 1 mM dEDTA at 25 °C.

For the real-time NMR experiments, 50 µM of ^15^N-labelled hDVL3 IDR2 was mixed with 1 µM CK1ε (aa 1-416) or 1 µM CK1δ (aa 1-415) in the buffer containing 50 mM KCl, 50 mM phosphate buffer [pH 6.5], 1mM dEDTA, 2 mM ATP and 10 mM MgCl_2_. The phosphorylation was initiated by the addition of CK1 isoform into the NMR tube and ^1^H-^15^N HSQC spectra were recorded every 30 minutes for 20 hours at 25 °C. Spectra were processed in TopSpin 4.0.6 and analysed in NMRFAM-SPARKY^69^ and Gnuplot 4.6.

For the analysis of real-time phosphorylation, peak intensity was measured for each unphosphorylated S/T or neighbouring residue in every spectrum and normalized to the reference spectrum at time zero. An exponential decay function was used to fit the decreasing intensity of the peaks over time:

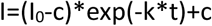

Where I is the peak intensity at time t, I_0_ is the peak intensity at time 0, c is the peak intensity at infinite time (level of phosphorylation), and k is the rate constant (phosphorylation rate). The fitting was done via k and c in Gnuplot 4.6.

The chemical shift perturbation (CSP) was calculated using the equation 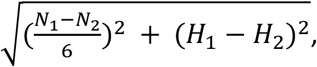, where N_1_ and H_1_ are the chemical shift values of the reference spectrum for nitrogen (N) and hydrogen (H), respectively, and N_2_ and H_2_ are the chemical shift values of N and H, respectively, in the titration series spectrum.

The CSP was subsequently used to calculate the dissociation constant (K_D_) using the following equation:

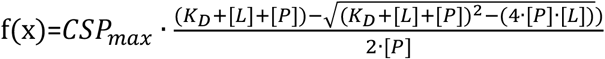

Where CSP_max_ the maximum value of CSP, [P] is the protein concentration, [L]=x represents the ligand concentration. The fitting was done via CSP_max_ and K_D_ in Gnuplot 4.6.

### HDX-MS

HDX-MS experiments were performed using a Synapt G2-Si HDMS coupled to nanoAQUITY UPLC with HDX Automation technology (Waters Corporation). hDVL3 PDZ-IDR2-DEP was concentrated to 34 µM, and optimization of the sequence coverage was performed on undeuterated controls. Analyses of phosphorylated and non-phosphorylated PDZ-IDR2-DEP were performed as follows: 3 µl of sample were diluted in 57 µl of either undeuterated (for the reference) or deuterated equilibration buffer (10 mM TrisCl [pH 7.5], 100 mM NaCl). The final percentage of deuterium in the deuterated buffer was 95%. Deuteration was performed at 20 °C for 0.5, 5, 10, 20, 40, 60, 120 and 240 min. Next, 50 µl of reaction sample was quenched in 50 µl of quench buffer (50 mM KH_2_PO_4_, 50mM K_2_HPO_4_ [pH 2.3]) at 0 °C. 80 µl of quenched sample was loaded onto a 50 µl loop and injected on an online Enzymate^TM^ pepsine column (Waters) maintained at 20 °C, with 0.2% formic acid at a flowrate of 100 µl/min. The peptides were then trapped at 0 °C on a Vanguard column (ACQUITY UPLC BEH C18 VanGuard Pre-column, 130 Å, 1.7 µm, 2.1 mm × 5 mm, Waters) for 3 min, before being loaded at 40 µl/min onto an Acquity UPLC column (ACQUITY UPLC BEH C18 Column, 1.7 µm, 1 mm × 100 mm, Waters) kept at 0 °C. Peptides were subsequently eluted with a linear gradient (0.2% formic acid in acetonitrile solvent at 5% up to 35% during the first 6 min, then up to 40% and 95% over 1 min each) and ionized directly by electrospray on a Synapt G2-Si mass spectrometer (Waters). HDMSE data were obtained by 20-30 V trap collision energy ramp. Lock mass accuracy correction was performed using a mixture of leucine enkephalin and GFP. Deuteration time points were performed in triplicates for each condition.

Peptide identification was performed from undeuterated data using ProteinLynx global Server (PLGS, version 3.0.3, Waters). Peptides were filtered by DynamX (version 3.0, Waters) using the following parameters: minimum intensity of 1,000, minimum product per amino acid of 0.2, maximum error for threshold of 10 ppm. All peptides identified in the IDR2 region were discarded and we only kept peptides present in both protein states, phosphorylated and nonphosphorylated, for comparison. All peptides were manually checked, and data was curated using DynamX. Back exchange was not corrected since we are measuring differential HDX and not absolute one. Statistical analysis of all ΔHDX data was performed using Deuteros 2.0^70^ and only peptides with a 99% confidence interval were considered.

### Charge calculation

The average charge calculations were done in R, version 4.2.2, using the idpr R package, version 1.8.0^71^. We used custom pKa dataset to include phosphorylated serine or threonine residues with pKa values of 5.6 or 5.9 for phosphorylated serine or phosphorylated threonine, respectively.^72^ For other residues, “IPC_peptide” pKa dataset was used.^73^ For the calculation of the local charge, we used the window of nine amino acid residues and pH = 6.5. Net charge was calculated at pH = 6.5 for *in vitro* experiments, and at pH = 7.2 for the cellular experiments, respectively.

For the intramolecular coupling plot, a sigmoidal curve was fitted using the equation: 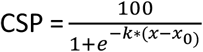, via k denoting the steepness of the sigmoidal function and via x_0_ denoting the inflection point of the curve (charge threshold). x is the negative net charge proximal to the DEP domain of phosphorylated IDR2. CSP was calculated as the weighted distance (see NMR spectroscopy) of the corresponding peak in the phosphorylated form from the WT peak in the non-phosphorylated form. CSPs of mutants were expressed as the percentage of WT CSP, the latter being 100 %. CSPs from T459, T480, K483, and S487 were averaged for WT, Δ S/T, and S/T-E. In the case of S/T-A, only CSPs from T459 and T480 were used as the other values reported on exchange phenomena (**Fig. S10**).

For the FZD association plots, a sigmoidal curve was fitted using the equation: 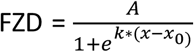, via k denoting the steepness of the sigmoidal function and via x_0_ denoting the inflection point of the curve (charge threshold). *A* is the maximum association. x is the negative net charge proximal to the DEP domain of phosphorylated IDR2. FZD was calculated as the Euclidean distance from the origin (0,0) in the volcano plot.

### Protein structures and visualization

All protein structure images were prepared in PyMOL.^74^ The electrostatics were calculated using the APBS Electrostatics,^75^ PyMol plugin. PDZ-IDR2-DEP structure was created in MODELLER 10.2.^76^ DEP-FZD structure was retrieved from PDB (8WMA).^11^ All other DVL structures were prepared using AlphaFold.^77,78^

### Site-directed mutagenesis and plasmids

Site-directed mutagenesis of pCDNA3.1-Flag-hDVL3^79^ or pETM11-hDVL3-243-496 to generate phospho-switch mutant series was performed by using of a QuikChange II XL site-directed mutagenesis kit (Agilent) according to the manufacturer’s instructions. The sequences of all mutants were verified by Sanger sequencing (Eurofins, Genomics). DNA encoding these mutants or WT DVL3 was later inserted into HA-TurboID-pCDNA5 vector using Prolonged Overlap Extension-PCR^80^.

pCDNA3.1-Flag-hDVL3 was used as WT hDVL3, pcDNA3-Ck1ε^81^ was used as WT CK1ε, pcDNA3.1-HA-hAxin^82^ was used as WT Axin1, and V5-TurboID-NES_pCDNA3^40^ was used as a source of TurboID tag (Addgene plasmid # 107169 ; http://n2t.net/addgene:107169 ; RRID:Addgene_107169).

### Cell transfection

For transfection, 200,000 cells/well were seeded in a 24-well plate. The next day, cells were transfected using polyethylenimine (PEI) at a concentration of 1 μg/ml and pH 7.4 with a PEI ratio of 6 μl of PEI/1 μg DNA. The plasmids and PEI mixture were separately diluted in plain DMEM (DMEM without FBS, and antibiotics), incubated at room temperature for 15 min, then mixed together in a total transfection volume of 50 μl/well. Samples were vortexed, centrifuged, and incubated for 20 min at room temperature before addition to the cells. After 6 hours, the medium containing the transfection mix was removed and exchanged for complete DMEM. For Western blotting (WB), cells were treated with Wnt-3a for 2 h, whereas for TOPflash the treatment was overnight. In rescue experiments, 0.01 μg DNA/well of hDVL3 was used, and the total amount of DNA was equalized to 0.4 μg DNA/well by pcDNA3.1 (Invitrogen).

### TOPflash assay

Cells for the dual-luciferase assay were transfected with hDVL3 plasmids as described above, along with 0.1 μg of the pRLtkLuc plasmid and 0.1 μg of the Super8X TOPflash plasmid per well. The dual-luciferase assay was performed using a Dual-Luciferase reporter assay system (E1960; Promega) according to the manufacturer’s instructions. Luminescence was measured using a Hidex Bioscan Plate Chameleon luminometer.

### Western blotting and Antibodies

Western blotting was performed according to standard procedures. Shortly, samples were resolved at 8% polyacrylamide (1068102, SERVA) gels by SDS-PAGE and transferred to Immobilon-P® PVDF membranes (IPVH00010, Merck). Membranes were blocked with 5% non-fat dried milk in the TBST buffer (10 mM Tris [pH 7.6], 120 mM NaCl, 0.08 % Tween) for 1 h at room temperature prior to the addition of primary antibodies.

Following antibodies were used for WB or immunocytofluorescence: DVL3 (sc-8027, Santa Cruz), β-Actin (4970, Cell Signaling), Flag M2 (F3165, Sigma-Aldrich), CK1ε (610446, BD Biosciences), HA.11 (901514, Biolegend).

### Immunocytofluorescence

200,000 cells/well (DVL1/DVL2/DVL3 triple KO HEK 293 T-REx) were seeded on a gelatin-coated coverslip in a 24-well plate. The cells were transfected the next day, as described above, along with 0.05 μg DNA/well of CK1ε or Axin1 plasmids when indicated. The following day after the transfection, cells were fixed in fresh 4% paraformaldehyde, permeabilized with 0.05% Triton X-100, blocked with PBTA (3% BSA, 0.25% Triton, 0.01% NaN_3_) for 1 h, and incubated overnight with primary antibodies. The next day, coverslips were washed in PBS and incubated with secondary antibodies conjugated with Alexa Fluor 488 (A21206; Invitrogen) and Alexa Fluor 568 (A-11004; Invitrogen). Nuclei were stained with 4′,6-diamidino-2-phenylindole (DAPI; D1306, Thermo Fischer Scientific). One hundred positive cells (N = 3, if not stated otherwise) were analysed per individual conditions using GraphPad Prism. Images were acquired with Leica SP8 system.

### TurboID

Cells were seeded 24-hours prior the experiment on 15-cm culture dishes. The expression of DVL was induced by 1 µg/ml doxycycline (HY-N0565B, MedChemExpress) treatment for 24 hours, and supplement with 50 µM biotin (SC204706A, Santa Cruz) afterwards. After 30 minutes, cells were washed with PBS and lysed with buffer containing 500 mM NaCl, 50 mM Tris [pH 7.4], 0.4% SDS, 2% Triton X-100, 1mM DTT and protease inhibitor cocktail for 15 minutes on ice. The lysed cells were sonicated twice for 10 seconds and incubated on ice for additional 15 minutes. The lysate was centrifuged at 16,500 g for 15 minutes at 4 °C, diluted 2 times with 50 mM Tris [pH 7.4] and incubated with Streptavidin Sepharose™ High Performance beads (17511301, Cytiva) while rotating for 18 hours at 4 °C. The next day, beads were washed two times with 2% (w/v) SDS, one time with the buffer containing 0.1% (w/v) deoxycholic acid, 1% (w/v) Triton X -100, 1mM EDTA, 500 mM NaCl, 50 mM HEPES [pH 7.5], one time with the buffer containing 0.5% (w/v) deoxycholic acid, 0.5% (w/v) NP-40, 1 mM EDTA, 250 mM LiCl, 10 mM Tris [pH 7.4] and two times with 50 mM Tris [pH 7.4].

## QUANTIFICATION AND STATISTICAL ANALYSIS

### Graphs and statistics

Graphs were prepared using R 4.2.2 if not stated otherwise. Graphs in figure S3A and S3B were prepared using Microsoft Excel 365. All the CSP plots were prepared using Gnuplot 4.6. The statistical significance of the phosphorylation of endogenous hDVL3 was tested by ANOVA in R 4.2.2. Statistical analysis of all ΔHDX data was performed using Hybrid statistical test in Deuteros 2.0.^70^ Statistical significance of TOPflash assay was tested by ANOVA in R 4.2.2. All the statistical information can be found in figure legends.

For the interactome analysis, the differential expression (DE) was tested by LIMMA (baits vs. TurboID tag, DVL3-mut vs. DVL3-wt). To remove non-specifically bound proteins, the DE results DVL3-mut vs. DVL3-wt were further filtered on all proteins enriched (adj. P Value < 0.05, log2 FC > 1) in any of the baits vs. TurboID tag. Volcano plots were created in R (v. 4.2.3) using packages ggplot2 (v. 3.5.0), ggrepel (v. 0.9.5) and ggprism (v. 1.0.5).

## Notes

### Competing Interest Statement

The authors have declared no competing interest.

