## Supplementary material for "A Wnt-induced conformational phospho-switch in DVL3 controls interaction with Frizzled receptors and Wnt/β-catenin signaling": All suplementary figures

### Figure S1

repetition 1

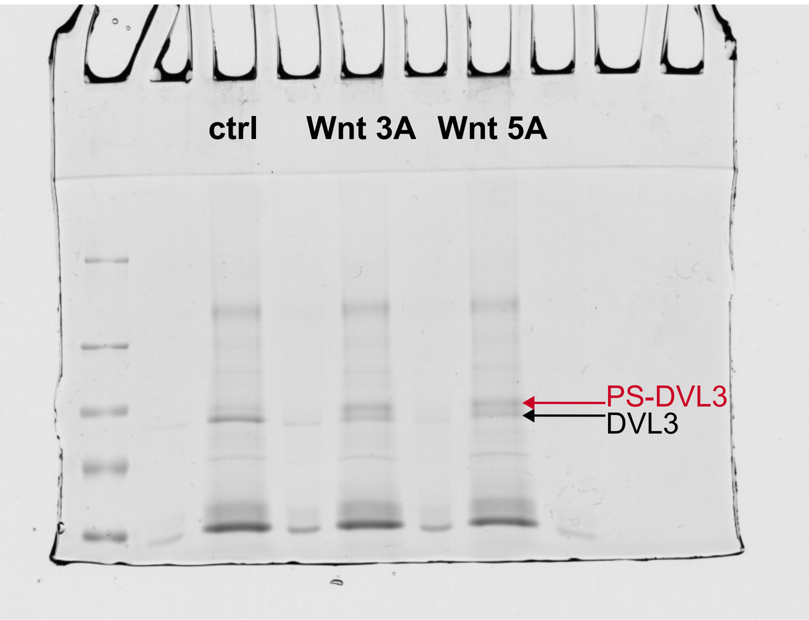


repetition 2

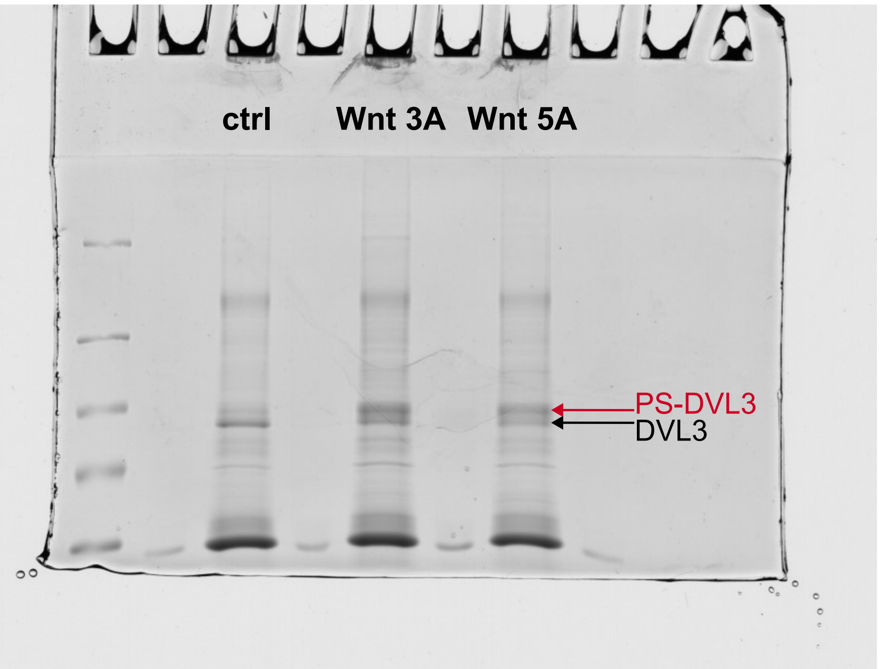


repetition 3

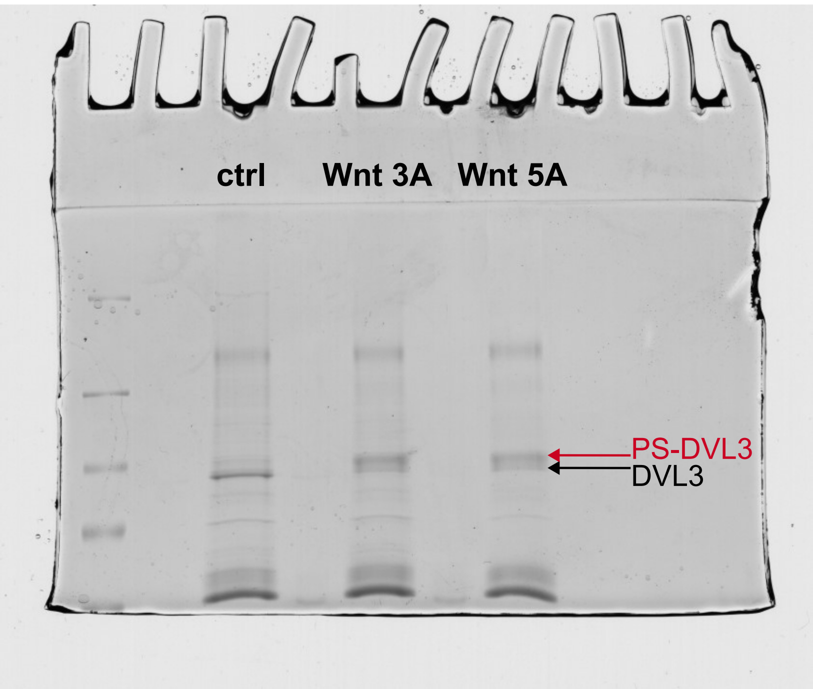

### Figure S2

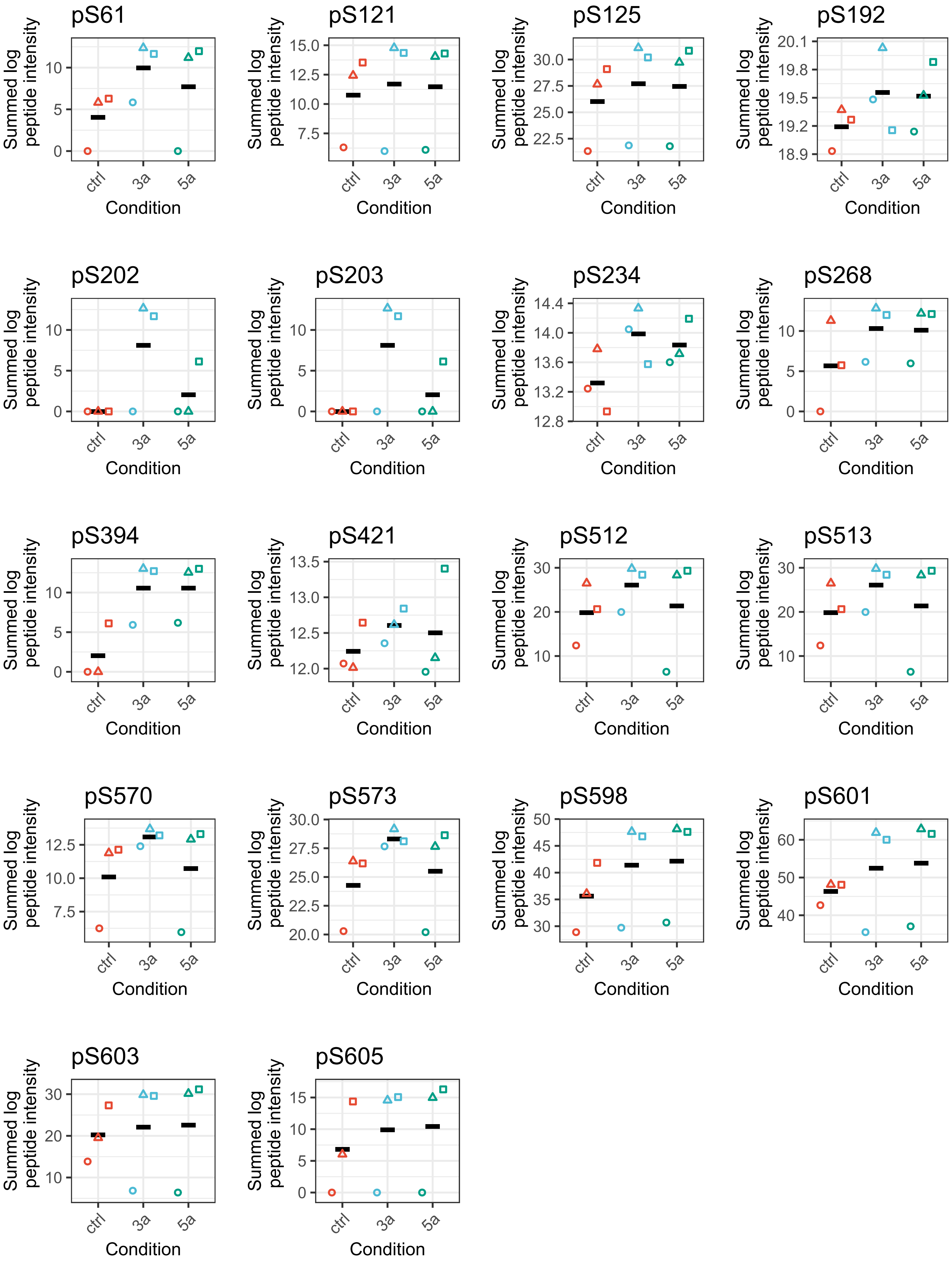

pS61
pS121
pS125
pS192
pS202
pS203
pS234
pS268
pS394
pS421
pS512
pS513
pS570
pS573
pS598
pS601
pS603
pS605
Summed log peptide intensity
Condition
ctrl
3a
5a

### Figure S3

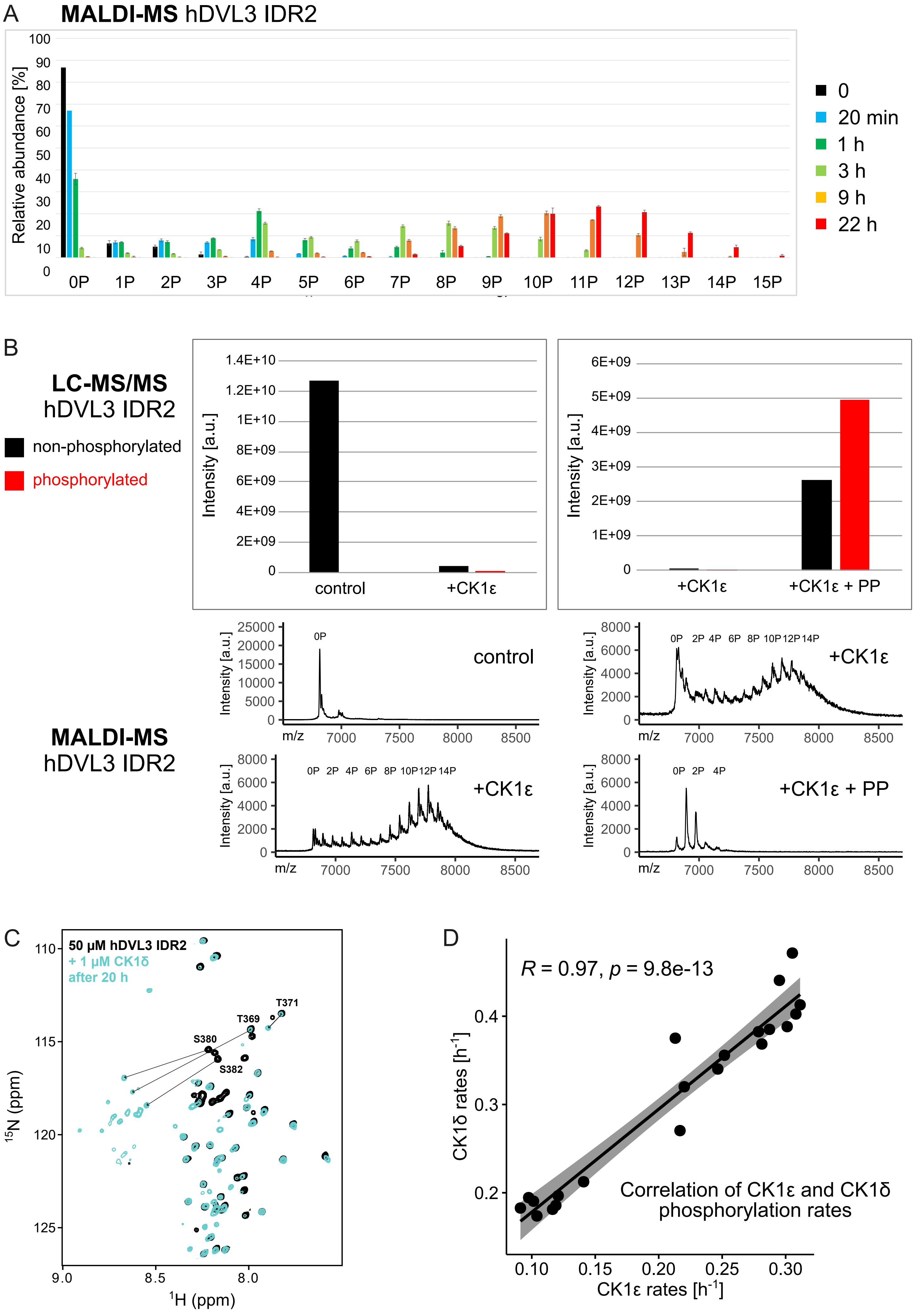
A
MALDI-MS hDVL3 IDR2
Relative abundance [%]
0P 1P 2P 3P 4P 5P 6P 7P 8P 9P 10P 11P 12P 13P 14P 15P
0
20 min
1 h
3 h
9 h
22 h
B
LC-MS/MS
hDVL3 IDR2
non-phosphorylated
phosphorylated
Intensity [a.u.]
control
+CK1ε
+CK1ε + PP
MALDI-MS
hDVL3 IDR2
m/z
0P 2P 4P 6P 8P 10P 12P 14P
C
50 µM hDVL3 IDR2
+ 1 µM CK1δ
after 20 h
T371
T369
S380
S382
15N (ppm)
1H (ppm)
D
R = 0.97, p = 9.8e-13
CK1δ rates [h-1]
CK1ε rates [h-1]
Correlation of CK1ε and CK1δ
phosphorylation rates

### Figure S4

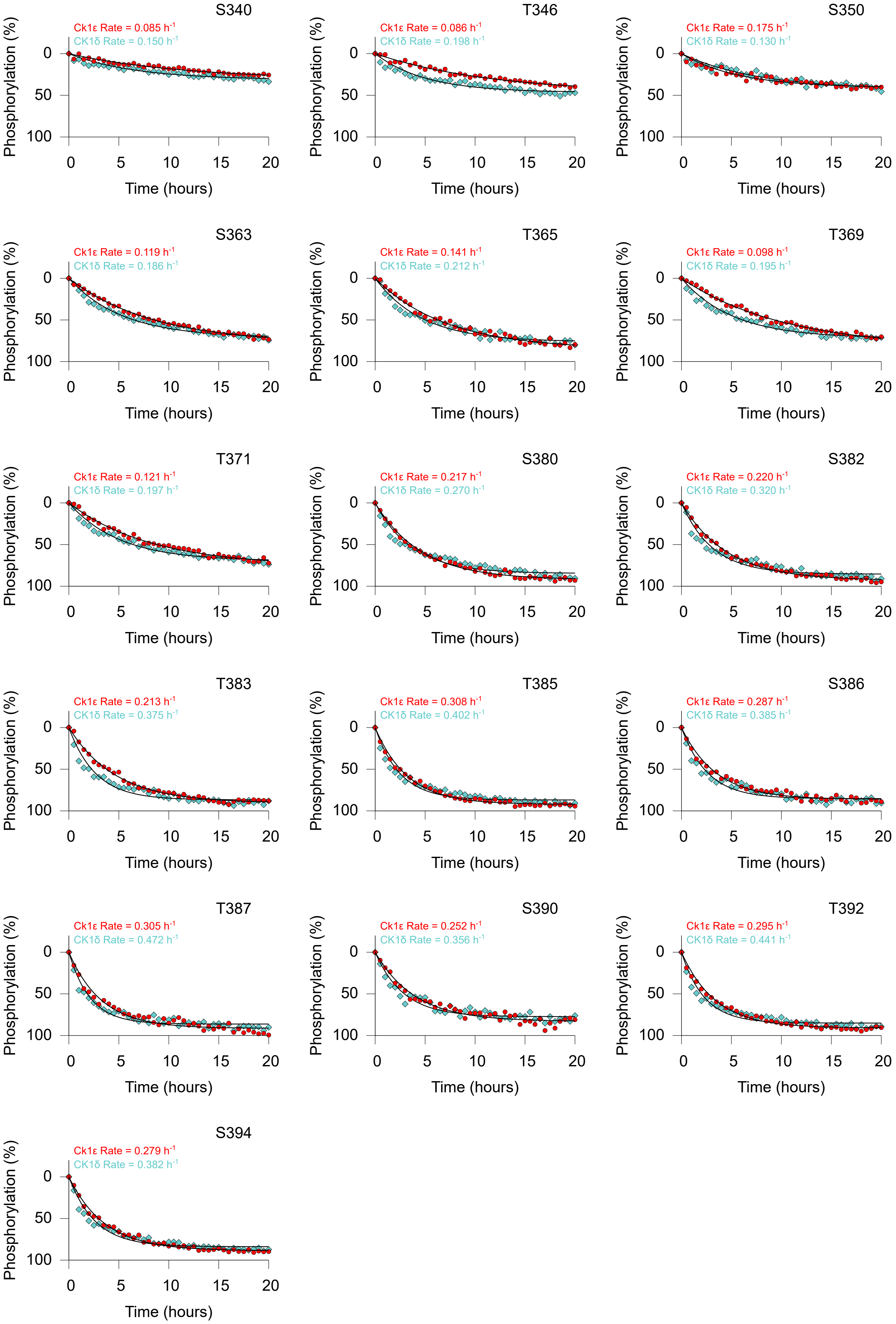
S340
Ck1ε Rate = 0.085 h-1
CK1δ Rate = 0.150 h-1
T346
Ck1ε Rate = 0.086 h-1
CK1δ Rate = 0.198 h-1
S350
Ck1ε Rate = 0.175 h-1
CK1δ Rate = 0.130 h-1
S363
Ck1ε Rate = 0.119 h-1
CK1δ Rate = 0.186 h-1
T365
Ck1ε Rate = 0.141 h-1
CK1δ Rate = 0.212 h-1
T369
Ck1ε Rate = 0.098 h-1
CK1δ Rate = 0.195 h-1
T371
Ck1ε Rate = 0.121 h-1
CK1δ Rate = 0.197 h-1
S380
Ck1ε Rate = 0.217 h-1
CK1δ Rate = 0.270 h-1
S382
Ck1ε Rate = 0.220 h-1
CK1δ Rate = 0.320 h-1
T383
Ck1ε Rate = 0.213 h-1
CK1δ Rate = 0.375 h-1
T385
Ck1ε Rate = 0.308 h-1
CK1δ Rate = 0.402 h-1
S386
Ck1ε Rate = 0.287 h-1
CK1δ Rate = 0.385 h-1
T387
Ck1ε Rate = 0.305 h-1
CK1δ Rate = 0.472 h-1
S390
Ck1ε Rate = 0.252 h-1
CK1δ Rate = 0.356 h-1
T392
Ck1ε Rate = 0.295 h-1
CK1δ Rate = 0.441 h-1
S394
Ck1ε Rate = 0.279 h-1
CK1δ Rate = 0.382 h-1
Phosphorylation (%)
Time (hours)

### Figure S5

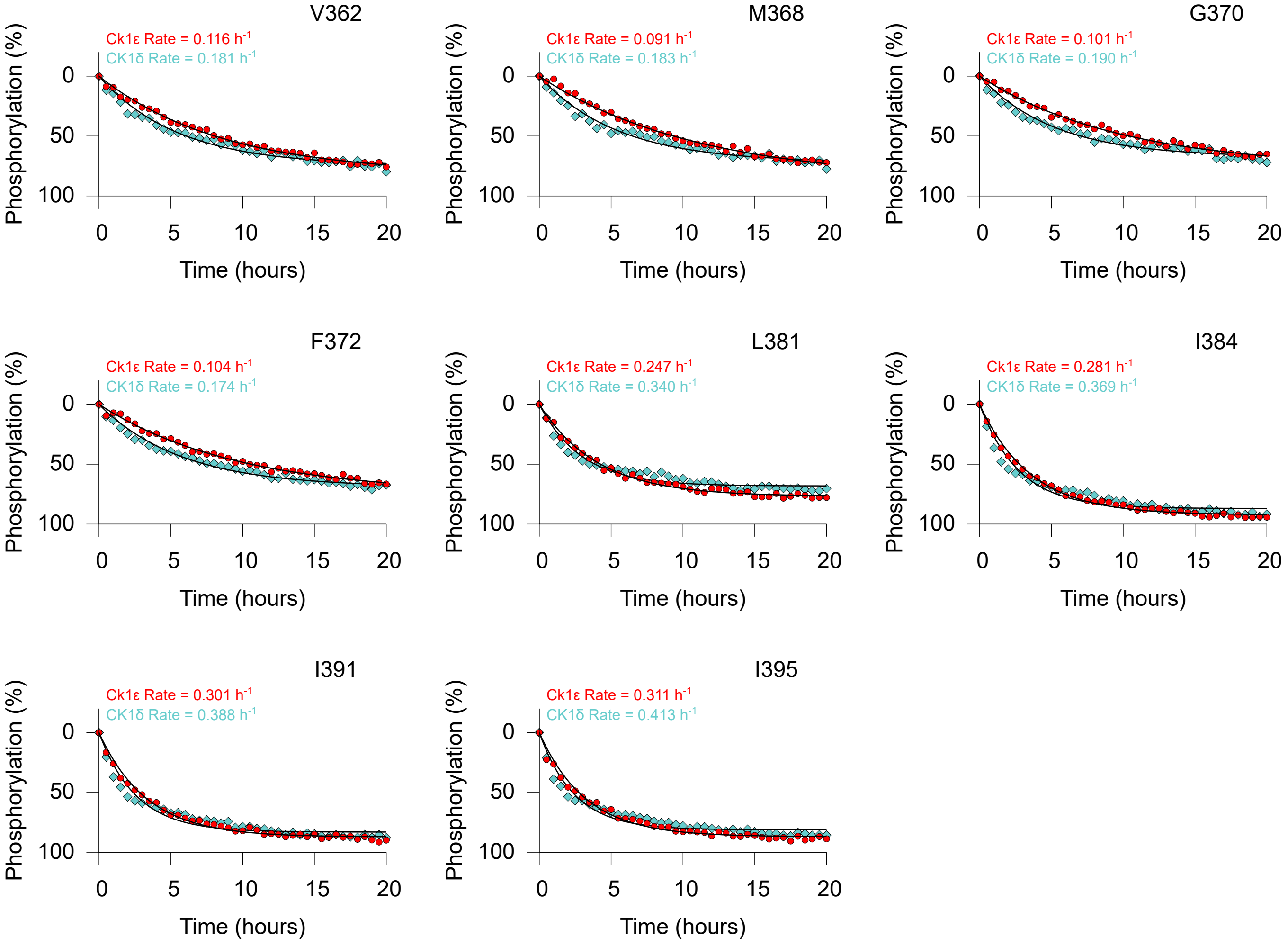
V362
Ck1ε Rate = 0.116 h-1
CK1δ Rate = 0.181 h-1
M368
Ck1ε Rate = 0.091 h-1
CK1δ Rate = 0.183 h-1
G370
Ck1ε Rate = 0.101 h-1
CK1δ Rate = 0.190 h-1
F372
Ck1ε Rate = 0.104 h-1
CK1δ Rate = 0.174 h-1
L381
Ck1ε Rate = 0.247 h-1
CK1δ Rate = 0.340 h-1
I384
Ck1ε Rate = 0.281 h-1
CK1δ Rate = 0.369 h-1
I391
Ck1ε Rate = 0.301 h-1
CK1δ Rate = 0.388 h-1
I395
Ck1ε Rate = 0.311 h-1
CK1δ Rate = 0.413 h-1
Phosphorylation (%)
Time (hours)

### Figure S6

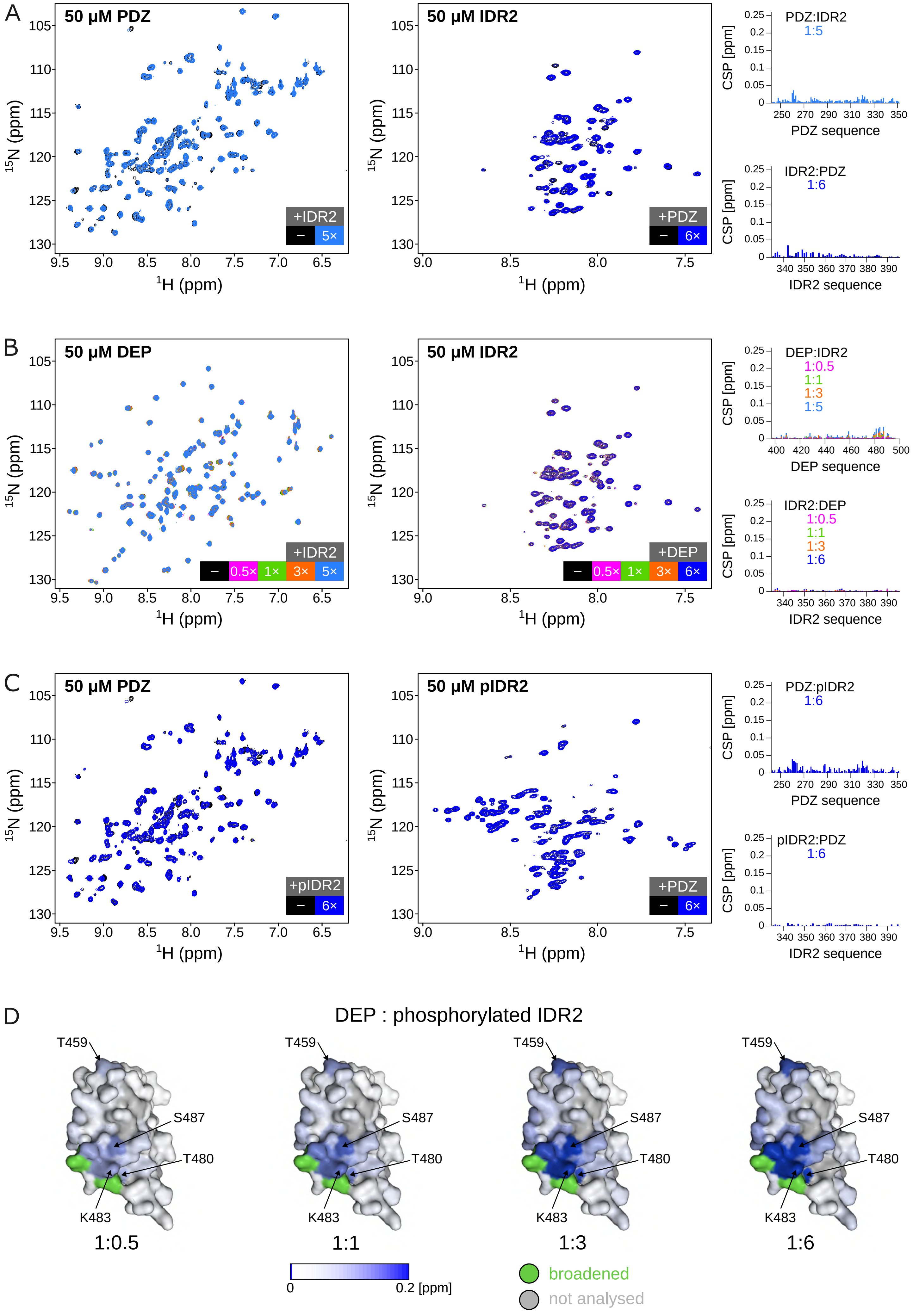

A
50 µM PDZ
+IDR2
–
5×
$^{15}$N (ppm)
$^{1}$H (ppm)
50 µM IDR2
+PDZ
–
6×
PDZ:IDR2
1:5
CSP [ppm]
PDZ sequence
IDR2:PDZ
1:6
IDR2 sequence
B
50 µM DEP
+IDR2
–
0.5×
1×
3×
5×
50 µM IDR2
+DEP
–
0.5×
1×
3×
6×
DEP:IDR2
1:0.5
1:1
1:3
1:5
DEP sequence
IDR2:DEP
1:0.5
1:1
1:3
1:6
C
50 µM PDZ
+pIDR2
–
6×
50 µM pIDR2
+PDZ
–
6×
PDZ:pIDR2
1:6
pIDR2:PDZ
1:6
D
DEP : phosphorylated IDR2
T459
S487
T480
K483
1:0.5
1:1
1:3
1:6
0
0.2 [ppm]
broadened
not analysed

### Figure S7

#### PDZ

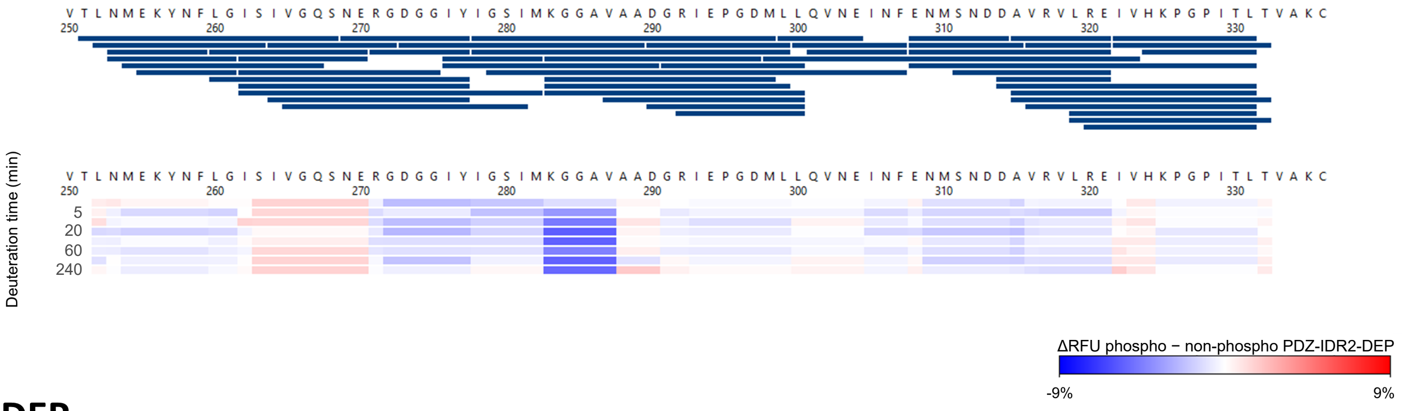


#### DEP

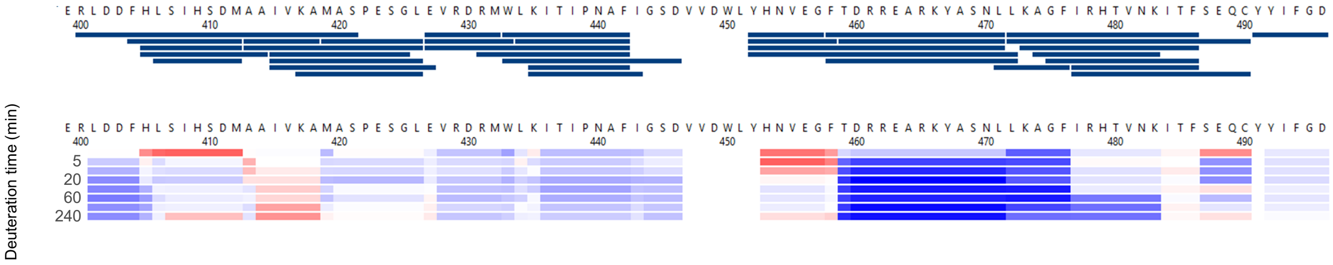

### Figure S8

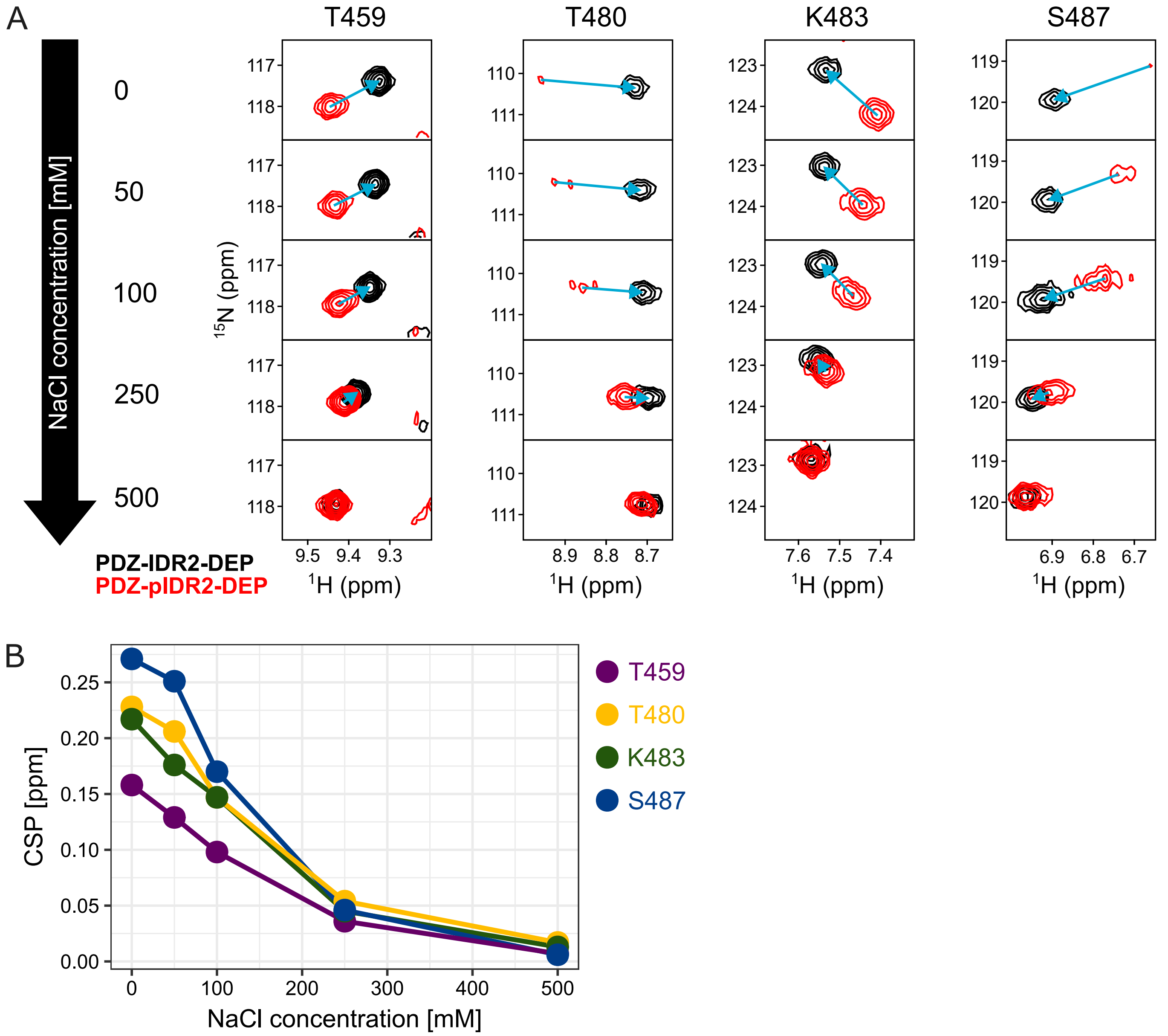

A
T459
T480
K483
S487
NaCl concentration [mM]
0
50
100
250
500
$^{15}$N (ppm)
$^{1}$H (ppm)
PDZ-IDR2-DEP
PDZ-pIDR2-DEP
B
CSP [ppm]
NaCl concentration [mM]
T459
T480
K483
S487

### Figure S9

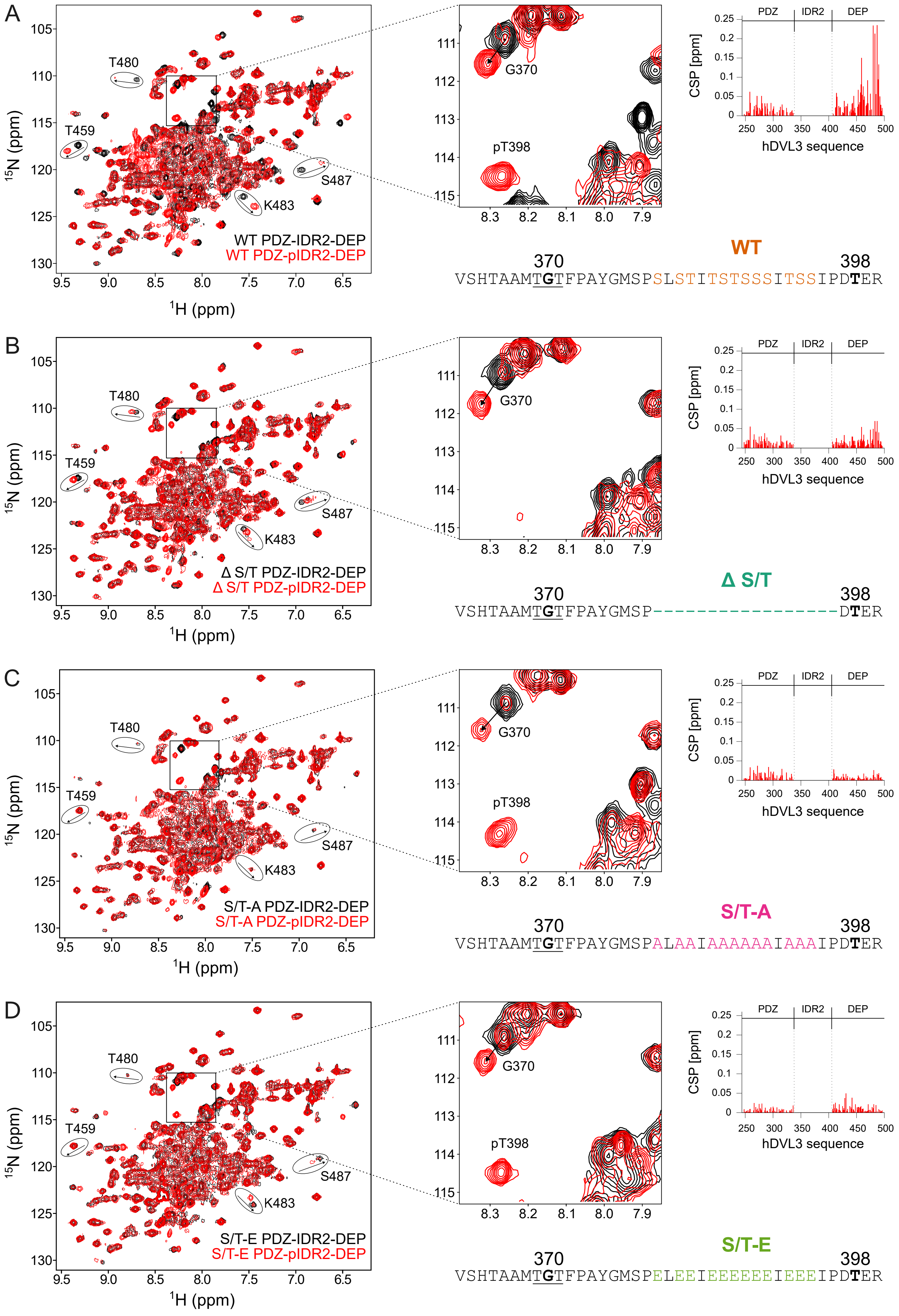

A
WT PDZ-IDR2-DEP
WT PDZ-pIDR2-DEP
T480
T459
S487
K483
G370
pT398
$^{15}$N (ppm)
$^{1}$H (ppm)
CSP [ppm]
PDZ
IDR2
DEP
hDVL3 sequence
WT
370
398
VSHTAAMTGTFPAYGMSPSLSTITSTSSSITSSIPDTER
B
Δ S/T PDZ-IDR2-DEP
Δ S/T PDZ-pIDR2-DEP
Δ S/T
VSHTAAMTGTFPAYGMSP----------------DTER
C
S/T-A PDZ-IDR2-DEP
S/T-A PDZ-pIDR2-DEP
S/T-A
VSHTAAMTGTFPAYGMSPALAAIAAAAAAIAAAIPDTER
D
S/T-E PDZ-IDR2-DEP
S/T-E PDZ-pIDR2-DEP
S/T-E
VSHTAAMTGTFPAYGMSPELEEIEEEEEEIEEEIPDTER

### Figure S10

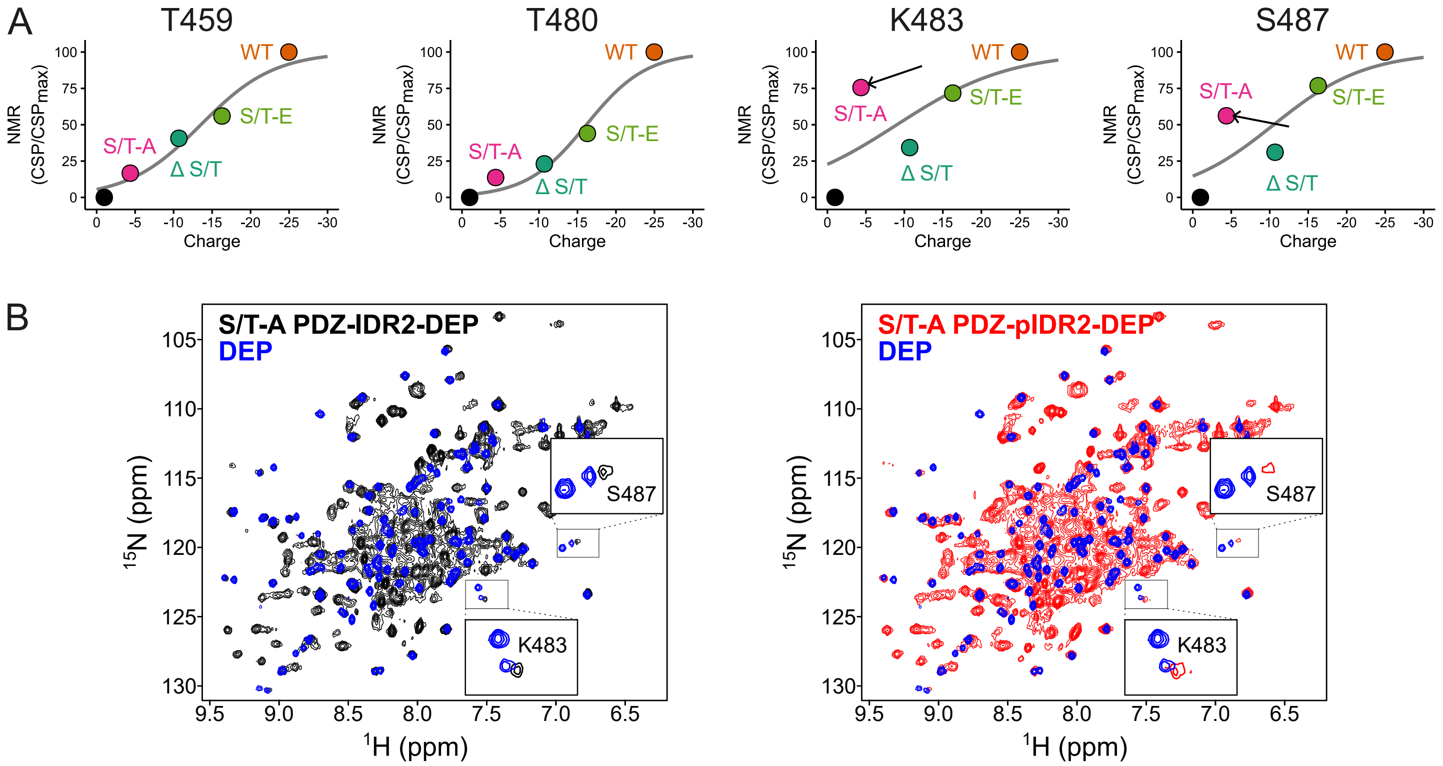

A
T459
T480
K483
S487
NMR (CSP/CSPmax)
Charge
WT
S/T-A
S/T-E
Δ S/T
B
S/T-A PDZ-IDR2-DEP
DEP
S/T-A PDZ-pIDR2-DEP
S487
K483
15N (ppm)
1H (ppm)

### Figure S11

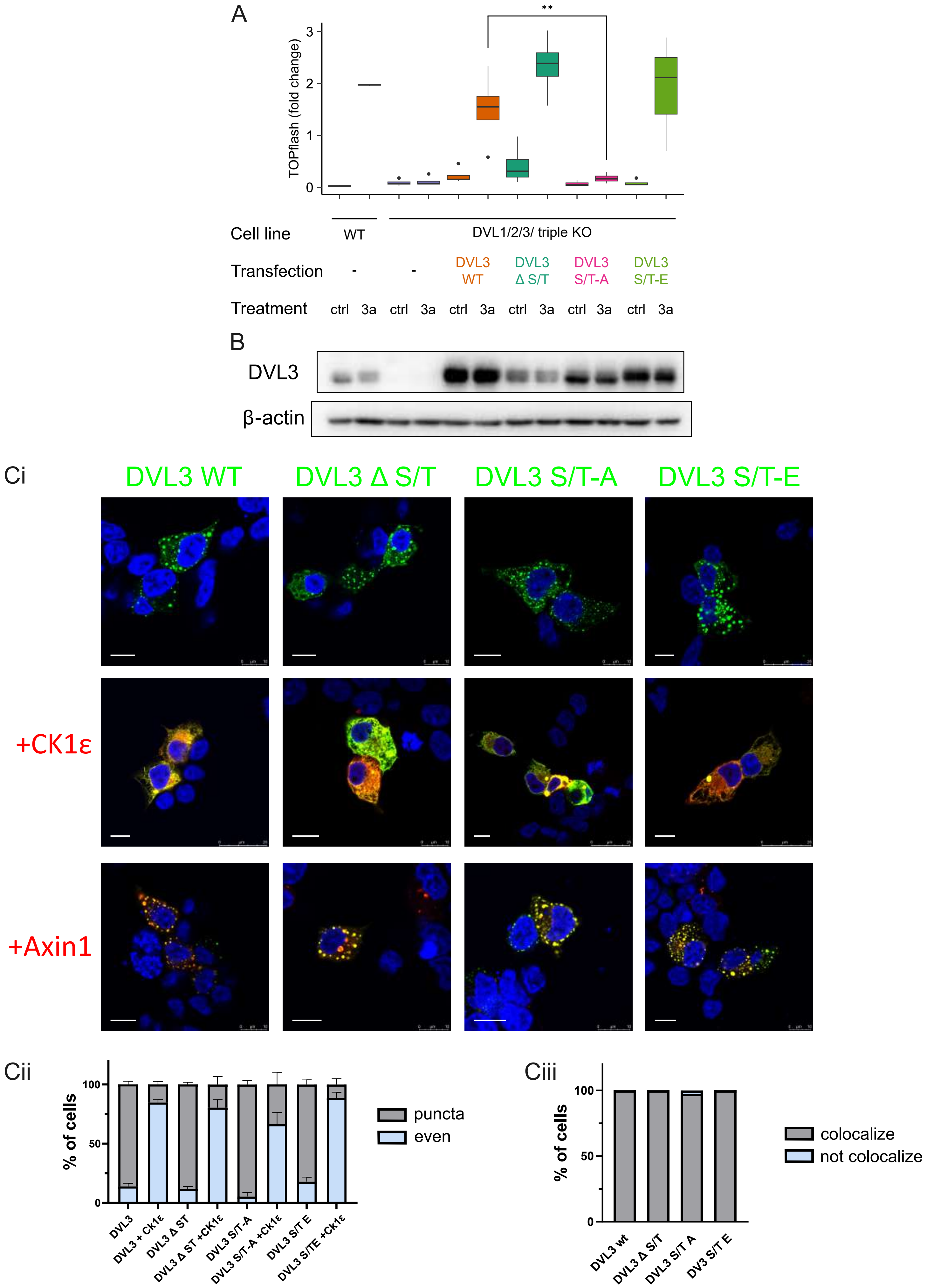
A
TOPflash (fold change)
**
Cell line
WT
DVL1/2/3/ triple KO
Transfection
DVL3 WT
DVL3 Δ S/T
DVL3 S/T-A
DVL3 S/T-E
Treatment
ctrl 3a ctrl 3a ctrl 3a ctrl 3a ctrl 3a ctrl 3a
B
DVL3
β-actin
Ci
DVL3 WT
DVL3 Δ S/T
DVL3 S/T-A
DVL3 S/T-E
+CK1ε
+Axin1
Cii
% of cells
puncta
even
DVL3
DVL3 + CK1ε
DVL3 Δ ST
DVL3 Δ ST +CK1ε
DVL3 S/T-A
DVL3 S/T-A +CK1ε
DVL3 S/T E
DVL3 S/TE +CK1ε
Ciii
% of cells
colocalize
not colocalize
DVL3 wt
DVL3 Δ S/T
DVL3 S/T A
DV3 S/T E

### Figure S12

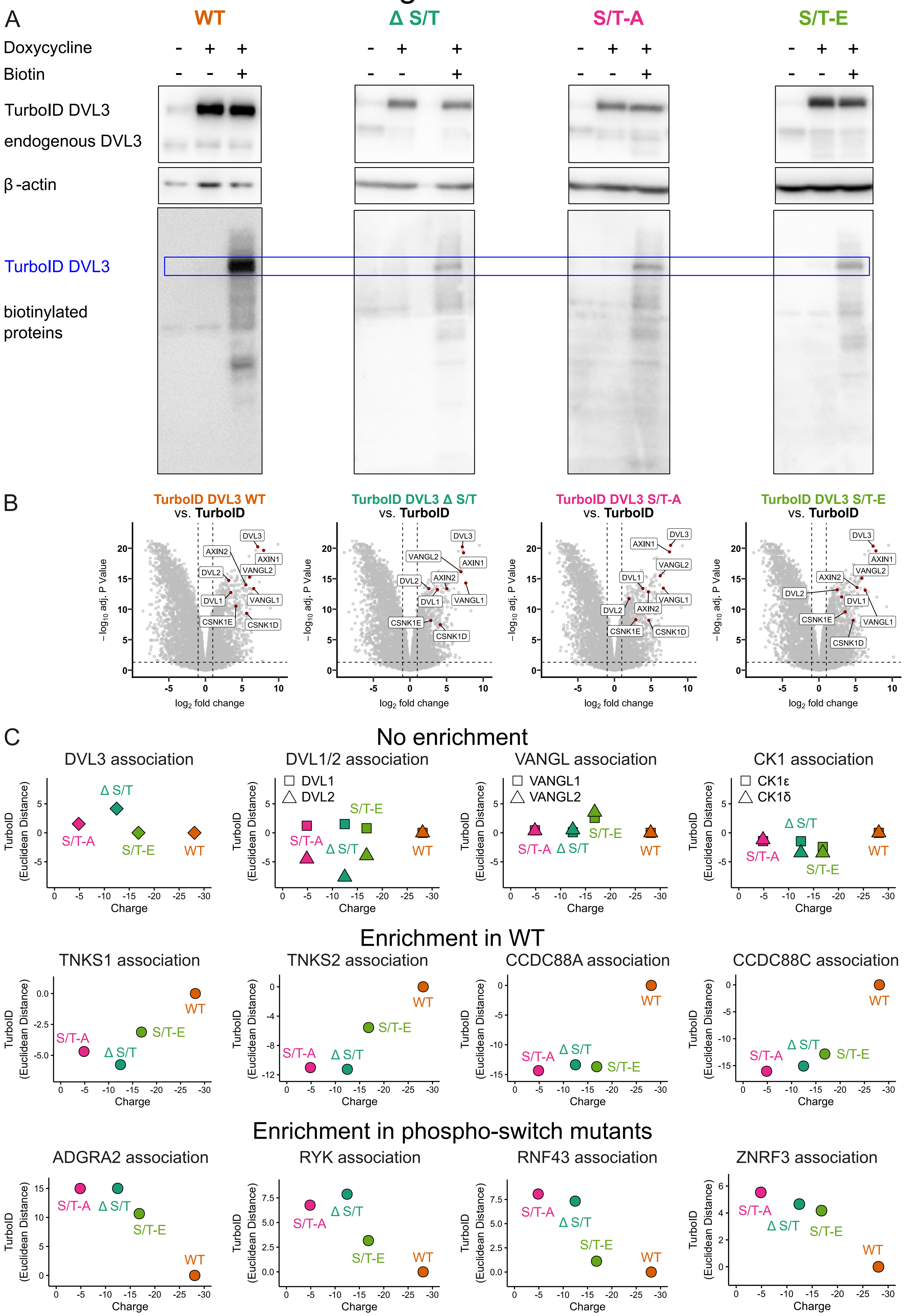
A
WT
Δ S/T
S/T-A
S/T-E
Doxycycline
Biotin
TurboID DVL3
endogenous DVL3
β -actin
TurboID DVL3
biotinylated proteins
B
TurboID DVL3 WT vs. TurboID
TurboID DVL3 Δ S/T vs. TurboID
TurboID DVL3 S/T-A vs. TurboID
TurboID DVL3 S/T-E vs. TurboID
DVL3
AXIN1
AXIN2
VANGL2
DVL2
DVL1
VANGL1
CSNK1E
CSNK1D
−log10 adj. P Value
log2 fold change
C
No enrichment
DVL3 association
DVL1/2 association
VANGL association
CK1 association
DVL1
DVL2
VANGL1
VANGL2
CK1ε
CK1δ
Enrichment in WT
TNKS1 association
TNKS2 association
CCDC88A association
CCDC88C association
Enrichment in phospho-switch mutants
ADGRA2 association
RYK association
RNF43 association
ZNRF3 association
TurboID (Euclidean Distance)
Charge

### Figure S13

A

```
hDVL3(aa335-400) KCWDPSPRGCFTLPRSEPIRPIDPAAWVSHTAAMTGTFPAYGMSPSLSTITSTSSSITSSIPDTER
hDVL3 Δ S/T      KCWDPSPRGCFTLPRSEPIRPIDPAAWVSHTAAMTGTFPAYGMSP-----------------DTER
dDSH(aa338-388)  KCWDPNPKGYFTIPRTEPVRPIDPGAWVAHTQALT-SH----DSI-IADI------AE---PIKER
```

B

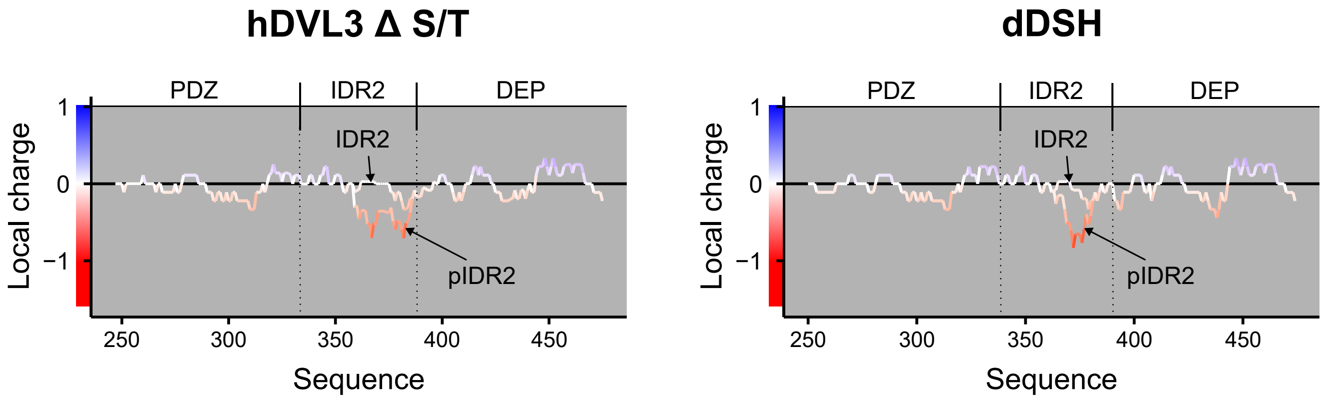
